# Human copy number variants are enriched in regions of low mappability

**DOI:** 10.1101/034165

**Authors:** Jean Monlong, Patrick Cossette, Caroline Meloche, Guy Rouleau, Simon L. Girard, Guillaume Bourque

## Abstract

Copy number variants (CNVs) are known to affect a large portion of the human genome and have been implicated in many diseases. Although whole-genome sequencing (WGS) can help identify CNVs, most analytical methods suffer from limited sensitivity and specificity, especially in regions of low mappability. To address this, we use `PopSV`, a CNV caller that relies on multiple samples to control for technical variation. We demonstrate that our calls are stable across different types of repeat-rich regions and validate the accuracy of our predictions using orthogonal approaches. Applying `PopSV` to 640 human genomes, we find that low-mappability regions are approximately 5 times more likely to harbor germline CNVs, in stark contrast to the nearly uniform distribution observed for somatic CNVs in 95 cancer genomes. In addition to known enrichments in segmental duplication and near centromeres and telomeres, we also report that CNVs are enriched in specific types of satellite and in some of the most recent families of transposable elements. Finally, using this comprehensive approach, we identify 3,455 regions with recurrent CNVs that were missing from existing catalogs. In particular, we identify 347 genes with a novel exonic CNV in low-mappability regions, including 29 genes previously associated with disease.

## 1 INTRODUCTION

Genomic variation of 50 base pairs or more are collectively known as structural variants (SVs) and can take several forms including deletions, duplications, novel insertions, translocations and inversions^1^. Copy number variants (CNVs) are unbalanced SVs, i.e. affecting DNA copy number, and include deletions and duplications. A wide range of mechanisms can produce SVs and is responsible for the diverse SV distribution across the genome, both in term of location and size^1,2,3^. In healthy individuals, SVs are estimated to cumulatively affect a higher proportion of the genome as compared to single nucleotide polymorphisms (SNPs)^4^. SVs have been associated with numerous diseases including Crohn’s Disease^5^, schizophrenia^6^, obesity^7^, epilepsy^8^, autism^9^, cancer^10^ and other inherited diseases^11,12^, and many SVs have a demonstrated detrimental effect.

While large SVs have been first studied using cytogenetic approaches and array-based technologies, whole-genome sequencing (WGS) is in theory capable of detecting SVs of any type and size^13^. Numerous methods have been implemented to detect SVs from WGS data using either paired-end information^14,15^, read-depth (RD) variation^16,17,18^, breakpoints detection through split-read approach ^19^ or de novo assembly^20^. CNVs, potentially the most impactful SVs, can be detected by any of these strategies but are often resolved with a RD approach as it directly looks for signs of copy number changes. However, several features of WGS experiments result in technical bias and continue to be a major challenge. For example, GC content^21^, mappability^22,23^, replication timing^24^, DNA quality and library preparation^25^ have a detrimental impact on the uniformity of the RD^26^. Unfortunately, this variability is difficult to fully correct for as it involves different factors, some of which are unknown, that vary from one experiment to another. This issue particularly impairs the detection of CNV with weaker signal, which is inevitable in regions of low-mappability that represent around 10% of the human genome^27^, for smaller CNVs or in cancer samples with cell heterogeneity or stromal contamination. As a result, existing approaches suffer from limited sensitivity and specificity^3,13^, especially in regions of low-complexity and low-mappability^22,23^. Even when problematic regions were masked and state-of-the-art bias correction^21,28^ were applied, we showed that technical variation in RD could still be found across three WGS datasets studied (Monlong et al., under review).

To control for technical variation, we recently developed a CNV detection method, `PopSV`, which uses a set of reference samples to detect abnormal RD (Monlong et al., under review). In each genome tested, the RD in a region is compared to the same region in the reference samples. `PopSV` differs from most previous RD methods, such as `RDXplorer`^29^ or `CNVnator`^17^, that scan the genome horizontally and look for regions that diverge from the expected global average. Even when approaches rely on a ratio between an aberrant sample and a control, such as `FREEC`^16^ or `BICseq`^30^, we showed that they do not sufficiently control for experiment-specific noise as compared to `PopSV` (Monlong et al., under review). Glusman et al.^31^ does go further and normalize the RD with pre-computed RD profiles that fit the GC-fingerprint of a sample but this approach excludes regions with extreme RD and does not integrate the variance observed in individual regions. `PopSV` is also different from approaches such as `cn.MOPS`^18^ and `Genome STRIP`^32^ that scan simultaneously the genome of several samples and fit a Bayesian or Gaussian mixture model in each region. Those methods have more power to detect CNVs present in several samples but may miss sample-specific events. Moreover, their basic normalization of the RD and fully parametric models forces them to conceal a sizable portion of the genome and variants with weaker signal. Finally, another strategy to improve the accuracy of CNV detection has been to use an ensemble approach that combines information from different methods relying on different types of reads. Large re-sequencing projects such as the 1000 Genome Project^3,33^ and the Genomes of Netherlands (GoNL) project^34,35^have adopted this strategy and have successfully identified many CNVs using an extensive panel of detection methods combined with low-throughput validation. Such a strategy increases the specificity of the calls at the cost of sensitivity.

Notably, with most of the tools and approaches described above, repeat-rich regions and other problematic regions of the genome are often removed or smoothed at some step of the analysis, to improve the accuracy of the calls. Although some methods^36,37^ try to model ambiguous mapping and repeat structure, only particular situations are addressed and, as a consequence, low-mappability regions are just scarcely covered in the most recent CNV catalogs^33^. This is unfortunate given that CNVs in such regions have already been associated with various diseases^12,38,39,40,41^ and that these regions are also more likely variable. Indeed, different types of genomic repeats are likely to contribute to CNV formation. For example, CNVs are known to be enriched in segmental duplications^2^ and short and long tandem repeats are also known to be highly polymorphic^42,43^. Moreover, repeat templates, like segmental duplications or transposable elements, can facilitate the formation of CNV through non-allelic homologous recombination and other mechanisms^44^.

Given these facts and the growing realization of the importance of repetitive regions in the genome^45,46^, we wanted to investigate the performance of `PopSV` in low-mappability regions and explore the comprehensive CNV distribution across a large cohort of healthy individuals. After showing that population-based RD measures are better than existing mappability estimates to correct for variable coverage, we apply `PopSV` to 640 WGS individuals from three human cohorts: a twin study with 45 individuals^47^, a renal cell carcinoma datasets with 95 tumor and control pairs^48^ and 500 unrelated individuals from the GoNL dataset^34^. We compare the performance of `PopSV` on these datasets with existing CNV detection methods in regions of low-mappability and validate the quality of the predictions across different repeat profiles using PCR validation. Additionally, using publicly available long-read sequencing data and assemblies, we show that `PopSV` is able to detect some highly ambiguous CNVs. Next, having demonstrated the quality of the `PopSV` calls, we characterize the patterns of CNVs across the human genome and produce a CNV catalog where variants of different types are better represented compared to existing catalogs. We further find that CNVs are significantly enriched in regions of low-mappability and in different classes of repeats. Finally, we identify novel CNV regions in low-mappability regions that were absent from previous CNV catalogs and describe their impact on protein-coding genes.

## 2 MATERIALS AND METHODS

### Data

Three publicly available WGS datasets were used. The first is a twin study^47^ with an average depth of 40x across 45 individuals, including 10 families of parents and monozygotic twins. The second is a renal cell carcinoma dataset^48^ (CageKid) with 95 tumor/normal pairs and an average depth of 54x. The third contains 500 unrelated individuals from the GoNL^34^ dataset with an average depth of 14x. In each study, the sequenced reads had been aligned using `bwa`^49^. See SUPPLEMENTARY INFORMATION for more details on access and read processing.

### Read count across the genome

The genome was fragmented in non-overlapping bins of fixed size. As a RD measure we used the number of properly mapped reads, defined as read pairs with correct orientation and insert size, and a mapping quality of 30 (Phred score) or more. In each sample, GC bias was corrected by fitting a LOESS model between the bin’s RD and the bin’s GC content. We used a bin size of 5 Kbp for most of the analysis. When specified, we used smaller bin sizes of 500 bp or 2 Kbp.

### RD and mappability estimates

To compare RD and mappability estimates in the Twin study, we first removed bins with extremely high RD if deviating from the median RD by more than 5 standard deviation. The RD across the different samples were then combined and quantile normalized. For each bin, we computed the average RD and standard deviation across the samples. We downloaded the mappability track for hg19^27^ and computed the average mappability in each bin. We compared the RD in one randomly selected sample with the mappability estimates and with the inter-sample RD average. To correct for the variation explained by the mappability estimates we fitted a generalized additive model using a cubic regression spline between the mappability estimates and RD in the sample (see SUPPLEMENTARY INFORMATION). With these estimations and the global standard deviation we computed a Z-score for each bin. A similar set of Z-scores was computed using the inter-sample average and standard deviation. The normality of these two Z-score distributions were compared in term of excess kurtosis and skewness. The Z-score distributions were also compared in different mappability intervals. Finally, 45 samples of each cohort were combined and their RD quantile normalized. The inter-sample RD mean and standard deviation were then computed separately in each cohort and compared with the mappability estimates and RD in the selected sample.

### PopSV approach for CNV detection

`PopSV` was first described and applied in a CNV analysis of epilepsy patients (Monlong et al., under review). Briefly, a set of samples are chosen as reference and used to guide the normalization of each bin. After normalization the average RD and standard deviation in each bin are saved and used to transform the RD in all samples into Z-scores. CNVs are called in each sample when the RD is significantly higher or lower than in the reference samples. The Z-scores can be segmented using the circular binary segmentation^50^ or after statistical testing at the bin level. More details are available in the original publication (Monlong et al., under review) and in the SUPPLEMENTARY INFORMATION. With `PopSV` there is no filtering, masking, smoothing or altering of repeat-rich regions: all the regions with properly mapped reads are analyzed.

### Coverage track and low-mappability regions

The average RD in the reference samples, a feature used during CNV calling, was used as a coverage track. Bins with a RD lower than 4 standard deviation from the median were classified as *low-mappability* (or *low coverage*). To highlight the most challenging region, we also defined *extremely low coverage* regions if the average RD was lower than 100 reads. We overlapped these regions with protein-coding genes and segmental duplications (see SUPPLEMENTARY INFORMATION), and computed the distance to the nearest centromere, telomere or assembly gap. We also counted the number of protein-coding genes overlapping at least one low-coverage region.

### CNV detection using other methods

`FREEC`^16^ and `CNVnator`^17^ were run on each sample separately starting from the BAM files and using the same bin size as for `PopSV` (5 Kbp). `cn.MOPS`^18^ was run on the same GC-corrected bin counts than for `PopSV` and samples from the same dataset were jointly analyzed. After retrieving split reads using `YAHA`^51^, `LUMPY`^52^ was run and we kept all the deletions, duplications and intra-chromosomal translocations larger than 300 bp. See SUPPLEMENTARY INFORMATION for more details.

### Clustering samples using the CNV calls

The similarity between two samples is defined by the amount of sequence called in both divided by the average amount of sequence called (see SUPPLEMENTARY INFORMATION). This distance is used for hierarchical clustering of the samples in the Twin study using different linkage criteria (*average, complete* and *Ward*). The clustering was performed using calls in regions with extremely low coverage (≤100 reads on average in the reference samples) only. The Rand index estimated the concordance between the clustering and the known pedigree, grouping the samples per family (see SUPPLEMENTARY INFORMATION).

### Replication in twins

For each twin and each method, a CNV call was defined as *replicated* if also found in the other monozygotic twin but in less than 50% of the population to remove systematic errors. The frequency was computed by counting samples with any overlapping CNVs. In order to avoid missing calls with borderline significance, we used slightly less confident calls for the second twin (see SUPPLEMENTARY INFORMATION). For each method, we computed the number and proportion of *replicated* calls per sample. We computed these metrics using all the calls, calls in low-mappability regions only, calls in segmental duplications, calls overlapping annotated repeats and calls overlapping annotated satellites, all using a minimum overlap of 90% of the call’s sequence. Finally, we computed the replication estimates for calls located at 1 Mbp or less from a centromere, telomere or assembly gap.

### Replication between paired normal and tumor samples

The same approach was applied in the renal cancer dataset. Here, *replicated* calls were found in a normal sample and its paired tumor but in less than 50% of the normal samples.

### Replication estimates and reliable regions

Using CNV calls found in less than 50% of the population, we defined as *reliable* a 10 Kbp region where more than 90% of the overlapping calls were *replicated* calls. We then compared the number and proportion of reliable regions for each method and in different types of region. As before, we compared regions overlapping low-mappability regions, segmental duplications, annotated repeats, satellites, or located at less than 1Mbp from a centromere, telomere or assembly gap.

### Experimental validation

A subset of variants in the Twin study were experimentally validated. First, we randomly selected one-copy and two-copy deletions, among small (~ 700 bp) and large (~ 4 Kbp) variants among the calls produced with 500 bp and 5 Kbp bins. The calls were visually inspected to design PCR primers (see SUPPLEMENTARY INFORMATION). We randomly selected 20 regions from those with available PCR primers. Next, we randomly selected deletions overlapping low-mappability regions and called in 6 samples or fewer. Because RD could not be used efficiently to fine-tune the breakpoints’ location, we retrieved the reads (and their pairs) mapping to the region and assembled them (see SUPPLEMENTARY INFORMATION). We randomly selected 17 regions from those with PCR primers. In addition to gel electrophoresis, the amplified DNA of some regions was sequenced by Sanger sequencing.

### Analysis of CEPH12878

High coverage PCR-free Illumina WGS data for 30 samples, including CEPH12878, was downloaded from the 1000 Genomes Project (1000GP)^33^ (see SUPPLEMENTARY INFORMATION). `PopSV` was run using 5 Kbp bins and all the samples as reference. Using the same coverage track as before we selected all deletions in CEPH12878 overlapping low-mappability regions (at least 90% of the call). We first looked for support in CEPH12878 assemblies that used Illumina short-read sequencing, BioNano Genomics genome maps and either single molecule sequencing from the Pacific Biosciences (PacBio) platform^53^ or 10X Genomics linked-read sequencing ^54^. For each selected deletion from `PopSV`, we aligned the flanking reference sequences to the assemblies using BLAST^55^ (see SUPPLEMENTARY INFORMATION). When both flanks could be mapped to a contig, we visually inspected MUMmer plots^56^ which either supported the deletion, the reference genome sequence or were too noisy to assess. We further annotated the selected calls if they overlapped with the deletions identified in Pendleton et al.^53^ over a minimum of 1 Kbp. Finally, we downloaded the corrected PacBio reads and built a local assembly and consensus around each selected `PopSV` deletion (see SUPPLEMENTARY INFORMATION). We visually inspected MUMmer plots of the assembled and consensus sequences to confirm the presence of the deletion.

### CNV catalog

We called CNVs separately in each cohort with `PopSV` using as reference samples the 45 samples in the Twin study, the normal samples in the cancer dataset and 200 samples in the GoNL dataset. For the Twin study and the renal cancer dataset, `PopSV` was run using 500 bp bins and 5 Kbp bins. Because of the lower sequencing depth, `PopSV` was run using 2 Kbp bins and 5 Kbp bins for the GoNL dataset. For each sample, calls from the 2 different runs were merged when consistent (see SUPPLEMENTARY INFORMATION). To compute the total number of calls, we collapsed calls with a reciprocal overlap higher than 50%. The amount of sequence affected in a genome is computed by merging all the variants in the cohort and counting the affected bases in the reference genome.

### Comparison with the 1000 Genomes Project SV catalog

Autosomal deletions, duplications and CNVs from the 1000GP SV catalog^33^ were downloaded (see SUPPLEMENTARY INFORMATION). To compare the amount of CNV with `PopSV`, we removed deletions smaller than 300 bp as well as variants with high frequency (> 80%). We compared CNV frequency between the 620 unrelated samples and a down-sampled set of 620 randomly selected individuals from the 1000GP SV catalog. The frequency was derived for all the nucleotide that overlaps at least one CNV as the proportion of individuals with a CNV in this locus. The frequency distribution was computed separately for the different CNV types.

### Comparison with CNV catalogs from long-read studies

The SV catalog from Chaisson et al.^57^ was downloaded and overlapped with the CNV catalogs from 1000GP and `PopSV` results on our 640 genomes. Here, the 1000GP catalog contained deletions, duplications and CNVs of any size and frequency. Using control regions and logistic regression we tested for an enrichment of variants in the SV catalog from Chaisson et al.^57^ (see SUPPLEMENTARY INFORMATION). The analysis was performed separately on deletions, duplications, low-mappability regions and extremely low-mappability regions. The same analysis was performed using the SV catalog from Pendleton et al.^53^.

### Novel CNV regions

Using the 620 unrelated individuals across the three cohorts, we selected CNVs present in more than 1% of the population (7 individuals or more) and not overlapping any CNV in the 1000GP catalog^33^. We used deletions, duplications and CNVs of any size and frequency from the 1000GP. Novel CNVs were collapsed into novel CNV regions, i.e. contiguous regions in which each base is overlapped by at least one novel CNV. The novel CNV regions were annotated using the low-mappability and extremely low-mappability tracks.

### Distance to centromere, telomere and assembly gaps

The centromeres, telomeres and assembly gaps (CTGs) were retrieved from the gap track in UCSC^58^. In chromosomes with missing telomere annotation, we defined the telomere as the 10 Kbp region at the ends of chromosome. The distance from each variant to the nearest CTG was computed and represented as a cumulative proportion. Because this distribution changes with the size of the variants, we sampled random regions in the genome with similar sizes and computed the same distance distribution (see SUPPLEMENTARY INFORMATION). Thanks to this null distribution we were able to see if variants were located closer/further to CTG than expected by chance.

### Enrichment in genomic features

We tested for CNV enrichment in different genomic features: genes, exons, low-mappability regions, segmental duplications, satellites, simple repeats and transposable elements. The different satellite families, frequent simple repeat motives, transposable element families and sub-families were also tested. For each sample, we computed a fold-enrichment as the fold change in proportion of regions overlapping a feature between CNV and control regions (see SUPPLEMENTARY INFORMATION). The significance was assessed using logistic regression on the CNV and control regions. To control for the enrichment in segmental duplications we used control regions with similar overlap profile (see SUPPLEMENTARY INFORMATION). We also added a variable representing the overlap with segmental duplications as a co-factor in the logistic regression model. When numerous tests were performed, e.g. satellite families, simple repeat motives, transposable element families or sub-families, the P-values were corrected for multiple testing using Benjamini-Hochberg procedure. Finally, for each CNV and control region, we computed the proportion of the region overlapped by satellites, simple repeats and transposable elements.

### Overlap with gene annotation

Exons of protein-coding genes and promoter regions (10 Kbp upstream of the transcription start site) were extracted from the Gencode annotation v19. We counted how many genes overlapped a CNV in the population when considering exons only, exons and promoter region, or gene body and promoter region. In addition, we computed these numbers using only genes associated with a disease in the OMIM database (Online Mendelian Inheritance in Man; http://omim.org/). These numbers were also computed for CNVs that overlapped more than 90% of various classes of repeats. For example, Satellite-CNVs are CNVs with more than 90% of their region annotated as satellites.

## 3 RESULTS

### 3.1 Modeling RD using population-based measures instead of mappability scores

When counting uniquely mapped reads, the mappability of a region is a major predictor of the observed RD. Theoretical mappability estimates^27^ strongly correlated with the RD in a sample but many regions with intermediate mappability diverged from the predicted levels of RD (Fig. S1a). By computing the average RD across the 45 samples from the Twin study in each 5 Kbp bin we found that this divergence is consistent across samples and not simply due to a high RD variance (Fig. 1a). These mappability estimates only approximate RD variation and cannot explain the RD profile in numerous regions. In contrast, population-based metrics more directly estimate the expected RD level (Fig. S1b). Similarly to what was done in Monlong et al. (under review) in high-mappability regions, we hypothesized that population-based estimates of RD mean and standard deviation could be used directly and help analyze regions with reduced RD. To test this hypothesis, Z-scores corrected by the mappability-based estimates were compared to Z-scores derived from both the inter-sample mean and standard deviation. The population-based Z-scores better followed a Normal distribution with an excess kurtosis of 0.2 and skewness of 0.004 compared to 29.4 and -2.284 respectively for mappability-adjusted Z-scores (Fig. 1b). The distribution of the population-based Z-scores was also more stable across the mappability spectrum (Fig. 1c). When comparing samples from the three different datasets, we noticed cohort-specific profiles in term of RD level and variance even though RD had been quantile normalized (Fig. S1c and S1d), suggesting that population-based estimates will be better at capturing subtle cohort-specific variation.

**Figure 1:**
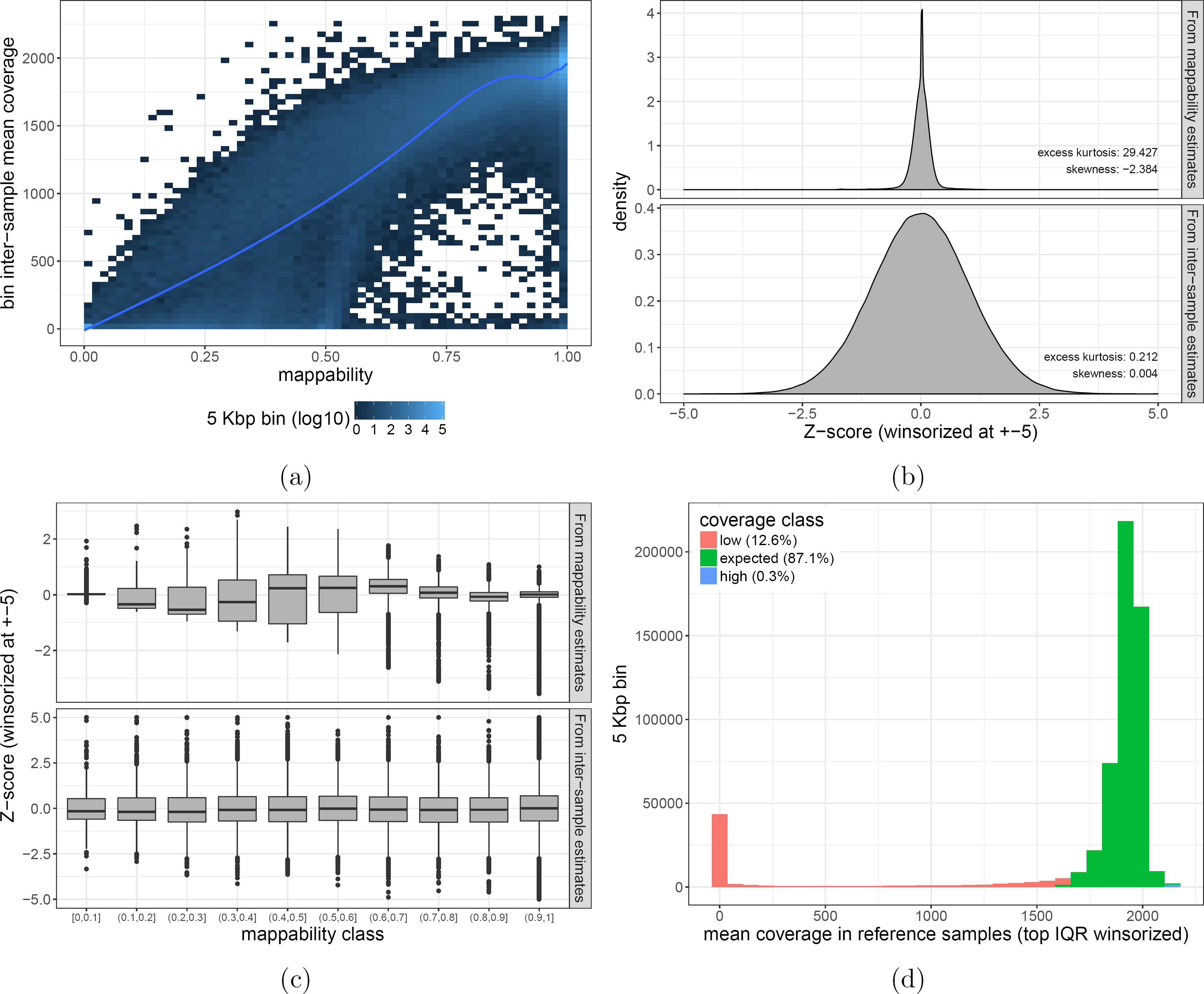
Mappability and population-based RD estimates. a) Inter-sample mean RD and average mappability in 5 Kbp bins. Regions with the same mappability estimate can have different RD levels. b) Z-score distribution. In *mappability*, Z-scores were computed from the mappability-predicted RD and global standard deviation; In *population estimates* from the inter-sample mean and standard deviation. c) Z-score distribution across the mappability spectrum. d) Average RD in the Twin study. The right-tail of the histogram was winsorized using the IQR and the different coverage classes are shown with colors.

These results suggest that a population-based strategy such as `PopSV` (Monlong et al., under review) could be extended to investigate CNVs in regions of low-mappability. To define low-mappability regions in the population, we used the average RD in the reference samples track produced by `PopSV`. In the Twin study for example, 12.6% of the covered 5 Kbp bins were labeled as low-coverage (Fig. 1d), more than half of which were regions with extremely low coverage (lower than 100 reads on average). Slightly fewer regions were labeled as low-coverage in the other cohorts (Fig. S2). As expected, low-coverage regions were depleted in gene content with only 15.3% of the 5 Kbp bins in these regions overlapping a protein-coding gene versus 48.8% for other regions. Nonetheless, 4,044 protein-coding genes overlapped a low-coverage region. Finally, 23.2% of the low-mappability regions overlapped segmental duplications and 69.1% were located at less than 1 Mbp from a centromere, telomere or assembly gap, versus respectively 2.9% and 8.8% for other regions.

### 3.2 Replication rates in regions of low-mappability

We previously demonstrated that CNV detection with `PopSV` was overall more sensitive than `FREEC`^16^, `CNVnator`^17^, `cn.MOPS`^18^ and `LUMPY`^52^ methods (Monlong et al., under review). In the following, we focused on the performance of `PopSV` in low-mappability regions. We first investigated the general concordance of the CNV calls with the pedigree in the Twin study. Using calls in extremely low-mappability regions (average RD below 100 reads) only, we clustered the individuals and compared the result to the known pedigree. We found that `PopSV` showed better concordance, as assessed by the Rand index (Fig. S3), compared to the other methods. Indeed, the clustering dendogram from `PopSV` calls, even in these challenging regions, captured almost perfectly the family relationships (Fig. 2a). We then investigated if the call replication rate was stable across different mappability profiles. Using calls present in less than 50% of the population to avoid systematic bias, the overall replication rate in the other twin was found to be 89.7%. Focusing on calls in low-coverage regions, we found a comparable replication rate of 92.5%. The replication rate remained constant in regions with different repeat profiles (Fig. 2b) such as regions overlapping segmental duplication, annotated repeats, or close to centromeres, telomeres and assembly gaps. In contrast, the other methods showed a reduced replication and higher variance in repeat-rich regions. The superior replication rate was complemented by a larger number of calls: `PopSV` called between 2.7 and 9.9 times more replicated CNVs per sample in low-coverage regions compared to the other methods. We observed the same results in the cancer dataset when comparing the agreement between germline events in normal/tumor pairs. `PopSV` had between 1.8 and 17.8 times more calls in low-mappability regions compared to the other methods and a stable replication rate across repeat profiles (Fig. S4). We next wanted to assess the performance in each region of the genome, rather than overall rates per sample, and used the replication in twins to identify regions with reliable calls. Again we observed that `PopSV` was as reliable overall as in regions with different repeat profiles (Fig. 2c). This analysis also showed that `PopSV` provides reliable calls in a larger fraction of the genome compared to other methods. The strongest gain was observed for regions overlapping satellites or overlapping almost completely annotated repeats, with around twice as many regions reliably called by `PopSV`. `cn.MOPS` showed the second best performance, especially in regions overlapping segmental duplications or close to centromeres, telomeres and assembly gap.

**Figure 2:**
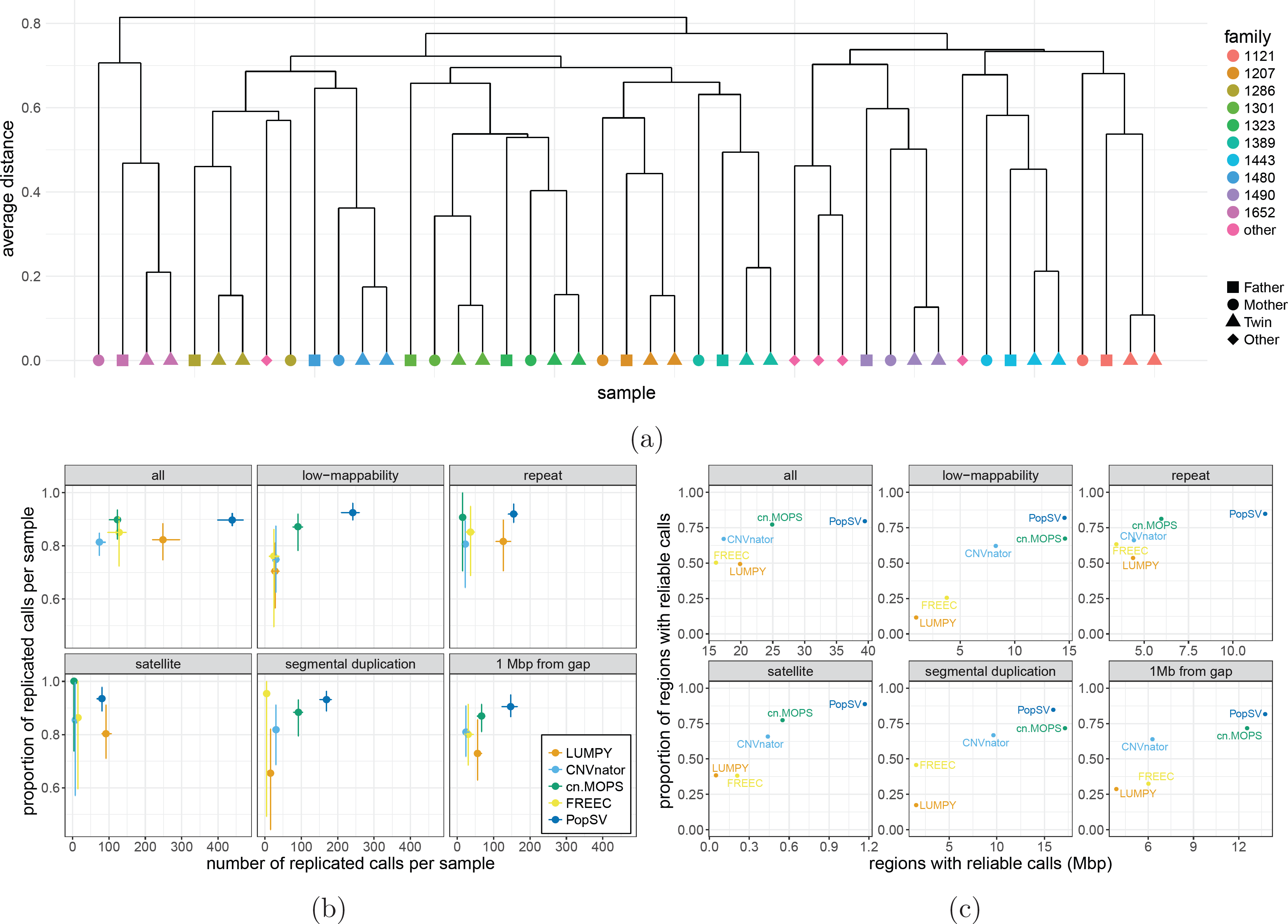
PopSV’s performance in low-mappability regions. a) Cluster using `PopSV` calls in extremely low coverage regions (below 100 reads). b) Proportion and number of calls replicated in the monozygotic twin. The point shows the median value per sample, the error bars the 95% confidence interval. c) Proportion and number of regions with reliable calls, computed from call replication in twins.

### 3.3 Validation of CNVs in regions of low-mappability

Using Real-Time PCR validation across 151 regions, we previously demonstrated that the replication estimates from the Twin dataset are consistent with experimental validation (Monlong et al., under review). We had tested variants of different types, sizes and frequencies and validated 90.7% of the calls, similar to our twin-based replication estimates. Here we tested additional deletions in individuals from the Twin study using PCR validation. We first validated randomly selected deletions and found a validation rate close to the overall replication rate, with 18 out of 20 deletions (90%) successfully validated (Table S1). In a second validation batch, we focused on rare deletions in low-mappability regions, of which 11 out of the 17 (65%) were successfully validated (Table S2). We noticed that the majority of the non-validated deletions were predicted to be smaller than 100 bp and most likely due to a problem during the breakpoint fine-tuning. If we consider only deletions larger than 100 bp, the validation rate in regions of low-mappability increased to 83% (10/12) once again close to `PopSV`’s replication rates in the Twin dataset.

Regions with extreme repeat content remained difficult to target and validate using PCR approaches. To further interrogate the performance of `PopSV` in those regions, we turned to whole-genome data from long-read sequencing technology. Publicly available assemblies for CEPH12878 samples confirmed several deletions called by `PopSV` in low-mappability regions. Out of the 14 homozygous deletions that could be assessed, 13 were confirmed in a contig, 12 of which were observed in both assemblies^53,54^. Only one region seemed to be a false positive, an assembled contig supporting the reference sequence in one assembly. Eleven regions could not be assessed because the flanks in the reference genome didn’t map to any assembled contigs or their MUMmer plots neither supported a deletion nor the reference sequence. In summary, we confirmed 92.8% of the homozygous deletions in low-mappability regions that could be compared with the assemblies. Deletions can be confirmed by direct comparison of the variant region and, if homozygous, should be present in the assembly. In contrast, heterozygous deletions could be missing from an assembly if only the reference allele was assembled. We confirmed 27 out of the 44 heterozygous deletions in low-mappability regions that could be assessed (Table S3). As expected, only one allele was supported for many regions: 16 regions with only the deleted allele observed and 17 regions with only the reference allele observed. Both deleted and reference alleles were observed for 11 variants. Although only 61.3% of the heterozygous deletion were confirmed, many variants might have been missed because of assembly preference to one allele, as suggested by the similar number of regions with only one supported allele. Using variants identified by Pendleton et al.^53^ and by assembling raw PacBio reads, we found support for 3 additional homozygous deletions and 15 heterozygous deletions that had remained inconclusive in the assembly comparison. Most of the regions that couldn’t be confirmed were located close to assembly gaps in the reference genome (Fig. S5). This observation highlighted that even with long-read sequencing data, it is not straightforward to clearly assess some genomic regions close to assembly gaps.

### 3.4 Global patterns of CNVs across the human genome

Having demonstrated the robustness of `PopSV` in low-mappability regions, we wanted to characterize the global patterns of CNVs across the human genome. We were especially interested in looking at calls in regions of low-mappability which represents between 9-12% of the human genome (Fig. 1d and S2). We started with an analysis of the twins and the normal samples in the renal cancer dataset, both of which have an average sequencing depth around 40X. `PopSV` was used to call CNV using 500 bp and 5 Kbp bins, which were then merged to create a final set of variants. On average per genome, 7.4 Mbp of the reference genome had abnormal read coverage, 4 Mbp showing an excess of reads indicating duplications and 3.4 Mbp showing a lack of reads indicating deletions (Table 1). In both datasets, the average variant size was around 3.7 Kbp and 70% of the variants found were smaller than 3 Kbp. We compared our numbers to equivalent CNVs detected in the most recent human SV catalog from the 1000 Genomes Project (1000GP), where 6.1 Mbp was found to be copy-number variable on average in each genome (Table S4). In those calls, we notice that no variants except for a few deletions were identified in regions of extremely low-mappability regions. Similarly, small duplications (< 3 Kbp) were absent from that catalog. In contrast, the set of variants identified by `PopSV` included variants in extremely low-mappability regions as well as small deletions and duplications (Table 1), explaining in part the ~ 20% increase in affected genome. While the study from the 1000GP^33^ explored a wider range of SVs, our catalog is likely more representative of the distribution of CNVs in a normal genome since a larger portion of the genome could be analyzed.

**Table 1:**
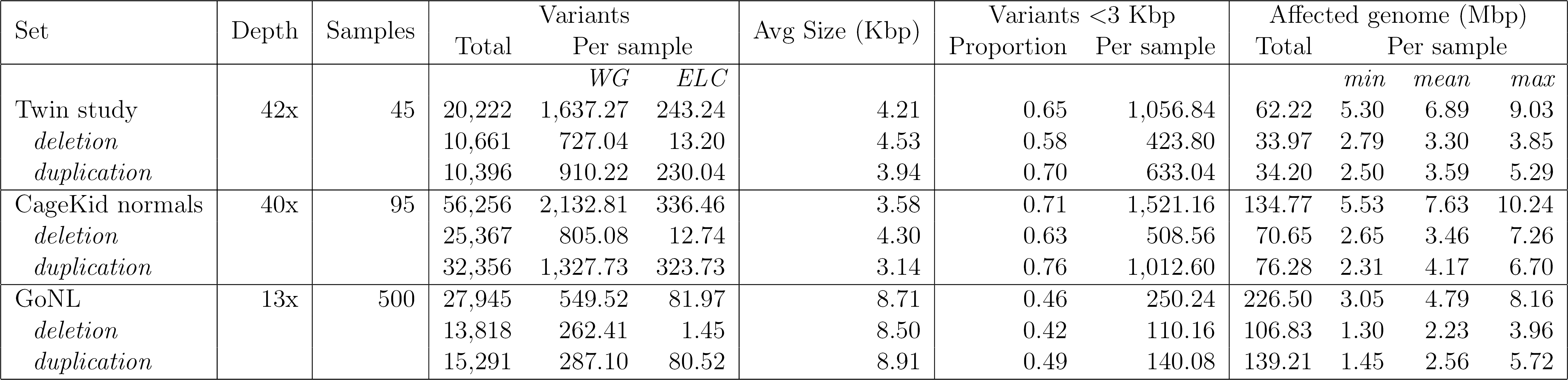
CNVs in the Twins, CageKid normals and GoNL datasets. WG: whole genome; ELC: extremely low-coverage regions. The *Total* number of variants is the total number after collapsing recurrent variants. *Affected genome* represents the amount of the reference genome that overlaps at least one CNV.

Next, we applied `PopSV` to the 500 unrelated samples from the GoNL cohort (Table 1). Due to a lower sequencing depth (~13X), we used bins of size 2 Kbp and 5Kbp, explaining the lower number of variants found in these samples. Nevertheless, a large sample size helps better characterize the frequency patterns and provides a more comprehensive map of rare CNVs. In total, across these three cohorts, 325.6 Mbp were found to be affected by a CNV with more duplications (50,856) detected than deletions (44,110). This contrasts with the CNVs reported by the 1000GP^33^ that were heavily skewed towards deletions (Table S4), likely due to the conservative ensemble approached used to detect CNVs. The frequency distribution of deletions and duplications found using `PopSV` were also much more balanced compared with the ones from the 1000GP^33^ (Fig. 3a).

**Figure 3:**
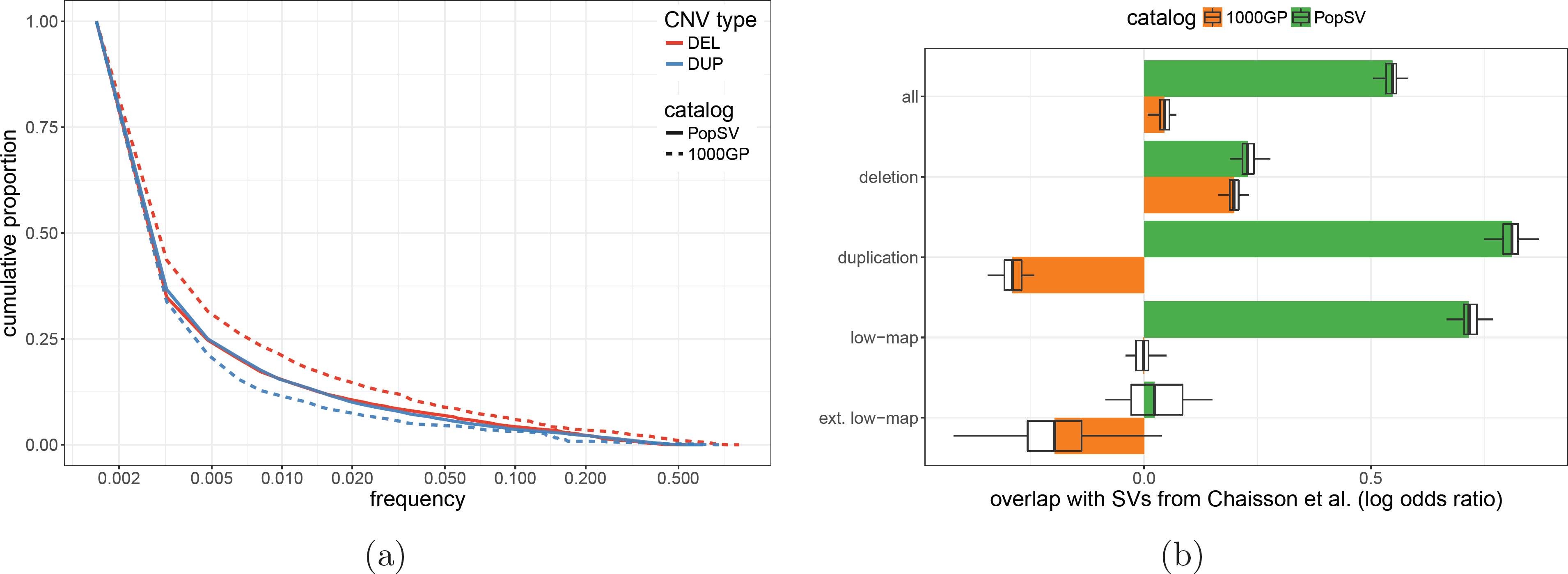
Comparison with CNV catalogs from the 1000 Genomes Project^33^ (1000GP) and a long-read sequencing study^57^. a) The x-axis represents the proportion of individuals with a CNV overlapping a region. The y-axis represents the cumulative proportion of the affected genome. b) Overlap with the SV catalog from Chaisson et al.^57^. In each cohort (color), the proportion of collapsed calls overlapping calls from Chaisson et al.^57^ or control regions with similar size distribution was modeled using a logistic regression. Boxplots show variation across 50 sampling of control regions. *low-map:* calls in low-mappability regions; *ext. low-map:* calls in extremely low-mappability regions.

We also compared our CNV catalog with an orthogonal set of calls from Chaisson et al.^57^ that were obtained using long-read sequencing. Although these calls came from a different genome, we expect both catalogs to share a number of common variants. We found a significant overlap between the two catalogs, overall and separately for deletions, duplications, low-mappability regions and extremely low-mappability regions (Fig 3b). In all categories, the overlap was stronger for `PopSV`’s catalog compared to the 1000GP CNV catalog. We noted that the enrichment for the 1000GP catalog disappeared for duplications and low-mappability regions but was even stronger for `PopSV`’s catalog. Like `PopSV`, the long-read sequencing study^57^ also found a better balance between deletions and duplications. Similar observations were made using another set of calls from long-read sequencing of the CEPH12878 sample^53^ (Fig. S6).

### 3.5 CNVs are enriched near centromeres and telomeres and in regions of low-mappability

Large CNVs have been shown to be enriched near centromeres, telomeres and assembly gaps (CTGs)^59^. We were interested in exploring this observation further using the set of high resolution calls from `PopSV`. We compared the distribution of CNVs calls made across the 3 datasets to randomly distributed regions of similar sizes (Fig. S7). In an average genome, we found that 33.5% of the CNVs calls were within 1 Mbp of a CTG, while we would have expected only 11.2% by chance. To verify that these observations were not simply a consequence of the methodology used, we also looked at the somatic CNVs (sCNVs) that we could detect in the renal cell carcinoma dataset. For this purpose, we extracted the variants found by `PopSV` in the tumor sample of an individual but missing from its paired normal sample. Reassuringly, and in contrast to germline CNVs, sCNVs were not preferentially found near CTGs (Fig. S7), with 11.1% of the sCNVs within 1 Mbp of a CTG.

After correcting for the distance to CTGs, we also observed a 4.7 fold-enrichment of variants in regions of low mappability (Fig. 4a). Segmental duplications (SD), DNA satellites and Short Tandem Repeats (STR) were also significantly enriched with fold-enrichment of 3.6, 2.6 and 1.2, respectively. The over-representation of CNVs in SDs has been described before^2^ and in a recent study^60^, half of the CNV base pairs were shown to overlap a SD. To investigate the contribution of low-mappability regions beyond SDs, we used matched control regions and included segmental duplication overlap in the logistic regression model. Even after controlling for this known enrichment, we found that CNVs overlapped low-coverage regions more than twice as much as expected (Fig. S8a). This two-fold enrichment is independent of the SD association and consistently observed in the 3 cohorts of normal genomes. In contrast to germline CNVs, sCNVs were once again found to be more uniformly distributed (Fig. 4a and S8a). These results suggest that the enrichments of germline CNVs near CTGs and in regions of low-mappability are unlikely to be the result of a methodological artifact.

**Figure 4:**
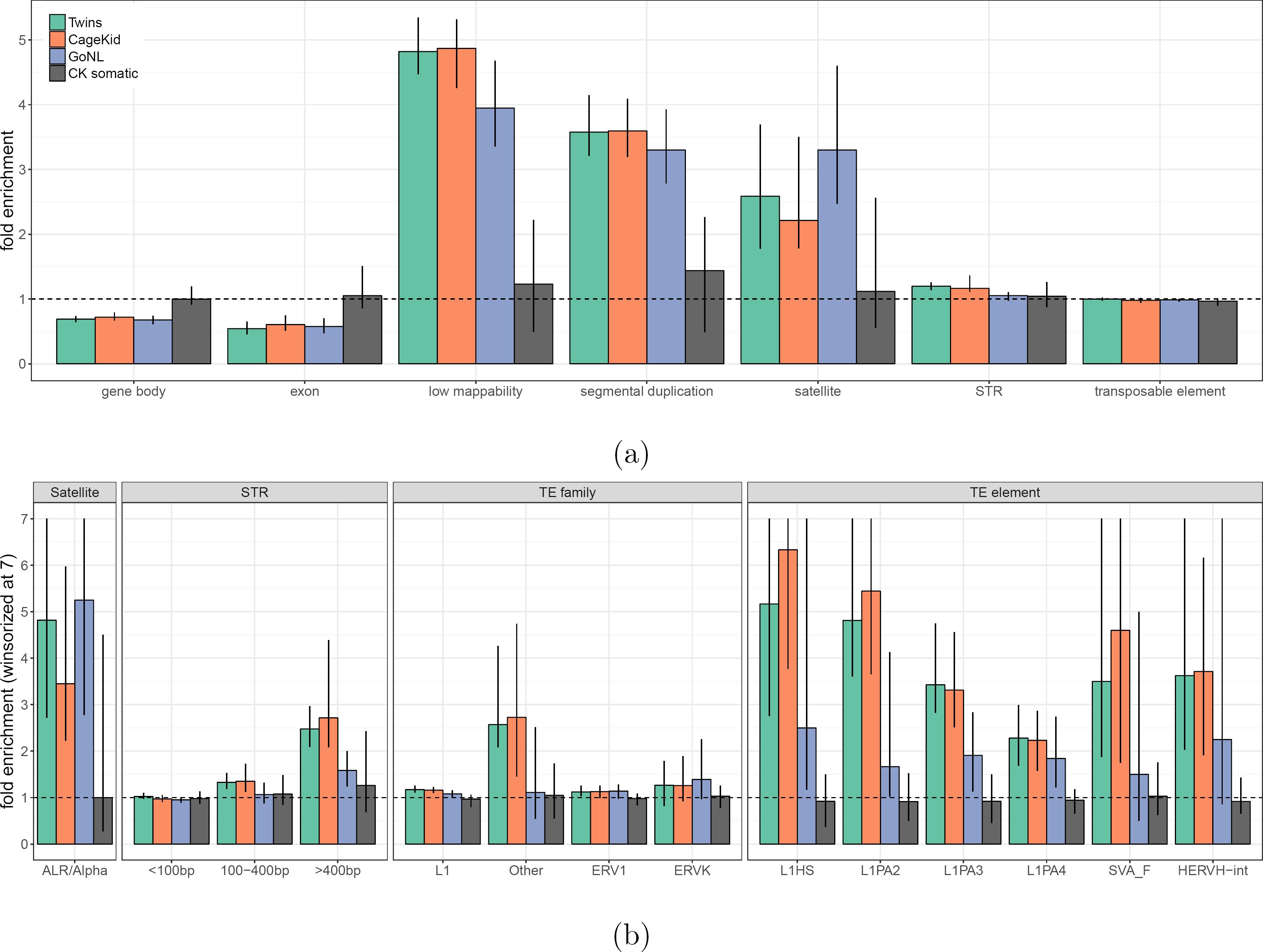
CNVs in normal genomes. a) Enrichment of CNVs in different genomic classes (x-axis) across different cohorts (colors) and controlling for the distance to centromere/telomere/gap. Bars show the median fold enrichment compared to control regions. The error bar represents 90% of the samples in the cohort. b) Enrichment of CNVs in repeat families (x-axis) controlling for the overlap with segmental duplication and distance to centromere/telomere/gap. The error bars were winsorized at 7 for clarity. *STR: Short Tandem Repeat; TE: Transposable Element.*

### 3.6 Various repeat families are more prone to harbor CNVs

We wanted to further characterize the distribution of germline CNVs in relation to different repeat classes and families. By comparing CNVs to the same control regions with matched overlap with SD and distance to CTGs we can look for patterns that are specific to repeat sub-families without the risk of being biased by the global enrichments (Fig. 4b). Using this approach, we found that CNVs were still significantly enriched in satellites repeats and in short tandem repeats (STRs) (P-value < 10^-4^, Fig. S8a), with fold-enrichments of 2.3 and 1.2 respectively.

Although it is known that DNA satellites and simple repeats are more unstable^61^, the extent to which CNVs are found in these regions in humans had, to our knowledge, not been systematically explored. Satellite repeats are grouped into distinct families depending on their repeated unit and we found that not all satellite repeats were equally likely to overlap a CNV (Fig. S8b). In particular, Alpha satellites have the highest and most significant enrichment (P-value < 10^-5^), with more than 3 times more CNVs than in the control regions (Fig. 4b). We noted that satellites tend to span completely CNVs (Fig. S9), suggesting that satellites are likely directly involved in the CNV formation. Short and long tandem repeats can be highly polymorphic^42,43^. Constrained by read length, recent studies^62,63^focused on variation of STRs smaller than 100 bp. In our analysis we found that CNVs were significantly enriched in the largest annotated STRs (>100 bp or >400 bp, Fig. 4b). STR can be grouped by motif and we further tested the largest and most frequent families (Fig. S8c). Except for the weak enrichment in *AT (TA)* repeats, the STR enrichment appeared mostly independent of the repeat motif. Here the repeats tend to overlap just a fraction of the variant, but a clear subset of the variants are fully covered by these tandem repeats (Fig. S9). Finally, although transposable elements (TEs) as a whole did not show enrichment (Fig. 4a), the “Other” repeat class, which contains SVA repeats, was found to be significantly enriched in the two higher depth datasets (Fig. 4b). Moreover, looking at TEs at the level of individual repeat families, we found a number of them to be significantly enriched including SVA F or L1Hs. Surprisingly, HERV-H, an older ERV sub-families, was also in the list of enriched TEs. This sub-family has been shown to be expressed and important in human embryonic stem cells^64,65^. Several families of older L1 repeats (e.g. L1PA2 to L1PA4) were also enriched and often implicated in what appears to be non-allelic homologous recombination (see examples in Fig. S10). Reassuringly, the somatic CNVs once again did not show any of these enrichments (Fig. 4b).

### 3.7 Impact of CNVs in regions of low-mappability

Compared to the latest 1000GP catalog^33^, we identified 3,455 novel regions with CNVs in more than 1% of the population. 81.3% of these regions were located in low-mappability regions while 18.4% were located in extremely low-mappability regions. Among the regions with a CNV in the CEPH12878 sample, we identified a deletion in the second intron of the *TRIM16* gene that was found by both Pendleton et al.^53^ and `PopSV`. Across the 640 individuals analyzed by `PopSV`, 12% carried the variant. Thanks to the long-read data, the exact breakpoints had been pinpointed in Pendleton et al.^53^ and it was in fact a SVA-F transposable element located within the 6 Kbp intron in the reference genome but absent from the assembled sequence. SVA-F is one of the youngest repeat family in the human genome and their high similarity remains a challenge for CNV analysis. Furthermore, the variant is located within a segmental duplication with 98.5% similarity and absent from public catalogs such as the 1000GP or GoNL. Another deletion supported by both public assemblies and local reassembly of the PacBio read was located 12 Kbp downstream of *TMPRSS11E.* 6.6% of the individuals carried the variant in the `PopSV` catalog. The assembled sequence helped pinpoint the breakpoints to an annotated L1PA2 in the reference genome. The variant was also located in a segmental duplication and absent from public catalogs such as the 1000GP or GoNL. Finally, a deletion affecting 8 different exons from the *CR1* gene was found by both Pendleton et al.^53^ and `PopSV` in CEPH12878. *CR1* has been associated with Alzheimer disease^66^ and is located within embedded segmental duplications with high similarity. The deletion was present in 3% of the population analyzed with `PopSV` but is absent from public CNV catalogs.

Overall, 7,206 protein-coding genes were found to have an exon overlapping a variant in at least one of the 640 normal genomes studied (Table 2). If we included the promoter regions (10 Kbp upstream of the transcription start site), at least 11,341 protein-coding genes were potentially affected by at least one CNV in the population. Focusing on regions of low-mappability, we found 4,285 different CNVs that were completely included in regions annotated as STR. These STR-CNVs overlapped the coding sequence of 45 protein-coding genes, and 286 genes when including the promoter region (Table 2). In contrast, for CNVs included in satellite regions, only 21 genes had an exon or the promoter region overlapping one of the 1,822 Satellite-CNVs. Finally, we focused on CNVs that were novel compared to the 1000GP^33^ and in low-mappability regions. Even there, 347 genes were found to have an exon overlapping such CNVs and this number increased to 560 when including the promoter regions. Out of these 347 genes, 29 were previously associated to a mendelian disorder in the OMIM database (Online Mendelian Inheritance in Man; http://omim.org/).

**Table 2:** Impact of CNVs on protein-coding genes. The *CNVs* number represents the number of different CNVs, after collapsing CNVs with more than 50% reciprocal overlap. Repeat CNV: more than 90% of the CNV is annotated as repeat. Genes are protein-coding genes and the promoter region is defined as the 10 Kbp region upstream of the transcription start site. *Novel CNVs* are located within regions annotated as novel compared to the 1000 Genome Project catalog.

| Set | CNVs | Genes with CNVs |  |  | OMIM genes with CNVs |  |  |
| --- | --- | --- | --- | --- | --- | --- | --- |
|  |  | Exon | + Promoter | + Intron | Exon | + Promoter | + Intron |
| *All CNVs* |  |  |  |  |  |  |  |
| All | 91,735 | 7,206 | 11,341 | 13,259 | 1,241 | 1,857 | 2,196 |
| Low coverage | 32,707 | 848 | 1,491 | 2,648 | 95 | 160 | 371 |
| Extremely low coverage | 9,348 | 304 | 401 | 442 | 11 | 14 | 25 |
| TE | 20,491 | 164 | 1,747 | 3,998 | 29 | 233 | 664 |
| STR | 4,285 | 45 | 286 | 748 | 5 | 39 | 129 |
| Satellite | 1,822 | 2 | 21 | 33 | 0 | 0 | 0 |
| *Novel CNVs* |  |  |  |  |  |  |  |
| All | 17,046 | 418 | 680 | 1,102 | 38 | 59 | 135 |
| Low coverage | 15,263 | 347 | 560 | 894 | 29 | 47 | 111 |
| Extremely low coverage | 6,591 | 189 | 263 | 285 | 5 | 6 | 8 |
| TE | 3,896 | 17 | 192 | 504 | 1 | 12 | 66 |
| STR | 1,806 | 14 | 81 | 230 | 0 | 9 | 41 |
| Satellite | 890 | 1 | 4 | 5 | 0 | 0 | 0 |

## 4 DISCUSSION

Despite the strong interest in CNVs because of their role in diseases, detecting them accurately has remained a challenge, especially in regions of low-mappability. This is mostly due to technical variation in RD that cannot be fully modeled by mappability estimates. Using a recently developed CNV-calling approach that relies on a set of reference samples to estimate the expected RD (Monlong et al., under review), we show that it is possible to accurately detect CNVs across the genome, even in repeat-rich regions. Indeed, using monozygotic twins and normal/tumor pairs, we were able to demonstrate that the performance of `PopSV` was stable and in most cases superior to other methods across different types of low-mappability regions. Although experimental validation can be challenging in these regions, we were able to confirm a number of deletions using PCR validation as well as variants in some of the most difficult regions by taking advantage of public datasets from long-read sequencing studies.

Notably, using `PopSV` on 140 normal genomes with high sequencing depth (~40X) and 500 additional samples with medium coverage (~13X), we found that regions of low mappability, which only represent ~10% of the genome, were around 5 times more likely to harbor CNVs. The fact that this enrichment was observed for germline events and not somatic events was both reassuring and interesting because of the implications on the selection forces at play. In particular, we were able for the first time to quantify the extent to which some regions in the genome are more prone to harbor such structural rearrangements. For instance, beyond the known enrichment in segmental duplications, we found genome-wide enrichments for different families of DNA satellites, simple repeats and TE, such as SVA, L1Hs and HERV-H. Moreover, although `PopSV` doesn’t fully characterize STR variation, it was able to detect CNVs in large annotated STRs. These CNVs could complement the output of STR detection methods that look for STR variation within sequencing reads and for this reason cannot test STRs longer than ~100 bp. Here, we found a strong CNV enrichment in STRs larger than 400 bp suggesting that large STRs should be included in genome-wide STR variation screens. Overall, having a more complete CNV catalog enabled an unbiased characterization of the CNV patterns across the genome and could potentially increase the power for trait-association studies.

Recent studies using long-read sequencing^57,53^found many novel SVs and highlighted variation involving complex repetitive DNA. The increased resolution and ability to span repeated regions expanded existing SV catalogs but only a handful of genomes have been sequenced in this way so far due to the higher cost of this technology. Although breakpoint and allele characterization is limited with short reads, we were able to detect the presence of such CNVs across a large population of normal genomes. Compared to previous studies, our CNV catalog strongly overlaps with the variants found by long-read sequencing studies in low-mappability regions. With hundreds of genomes at our disposal we identified frequent CNVs in repeat-rich regions that had escaped previous population-scale surveys. In the CEPH12878 sample, we independently identified low-mappability variants and showed that some novel deletions were recurrent in our cohort. For example, an exonic deletion in the *CR1* gene absent from public CNV catalogs was identified by the long-read sequencing and found in ~3% of the samples tested by `PopSV`. *CR1* has been associated with Alzheimer Disease^66^ thus this exonic deletion in a low-mappability region might be relevant for association studies. Using our full CNV catalog, we identified 3,455 novel regions that were not present in 1000G public SV database^33^ but found in more than 1% of our 640 genomes. These regions overlapped exons of 418 protein-coding genes, 38 of which were associated with a disease phenotype in the OMIM database. The amount of genes hit by CNVs in novel or low-mappability regions and the enrichment of CNVs in repeat-rich regions suggest that they be included in genome-wide surveys. As other types of variant are likely enriched in repeat-rich regions, we anticipate that population-based methods, such as `PopSV`, will facilitate the identification not only of CNVs but also of other types of SVs in both normal and cancer genomes.

## 5 DATA AND CODE AVAILABILITY

The `PopSV` R package and documentation are available at http://jmonlong.github.io/PopSV/. The scripts and instructions to reproduce the graphs and numbers in this study have been deposited at http://github.com/jmonlong/reppopsv/ and archived in https://doi.org/10.5281/zenodo.1181852.

## 6 ACESSION NUMBERS

The CNV catalog and annotations were deposited at https://figshare.com/s/8fd3007ebb0fbad09b6d. The raw sequences of the different datasets had already been deposited by their respective consortium (see SUPPLEMENTARY INFORMATION).

## 7 ACKNOWLEDGMENTS

We are grateful to the team of the Québec Study of Newborn twins who provided the twin dataset and the Cagekid consortium who provided the renal cancer dataset. This study also made use of data generated by the Genome of the Netherlands Project. A full list of the investigators is available from www.nlgenome.nl. Funding for the project was provided by the Netherlands Organization for Scientific Research under award number 184021007, dated July 9, 2009 and made available as a Rainbow Project of the Biobanking and Biomolecular Research Infrastructure Netherlands (BBMRI-NL). The sequencing was carried out in collaboration with the Beijing Institute for Genomics (BGI). Finally, we would like to thank Simon Gravel, Mathieu Blanchette, Mathieu Bourgey and Toby Dylan Hocking for helpful discussions.

## 9 SUPPLEMENTARY TABLES

**Table S1:**
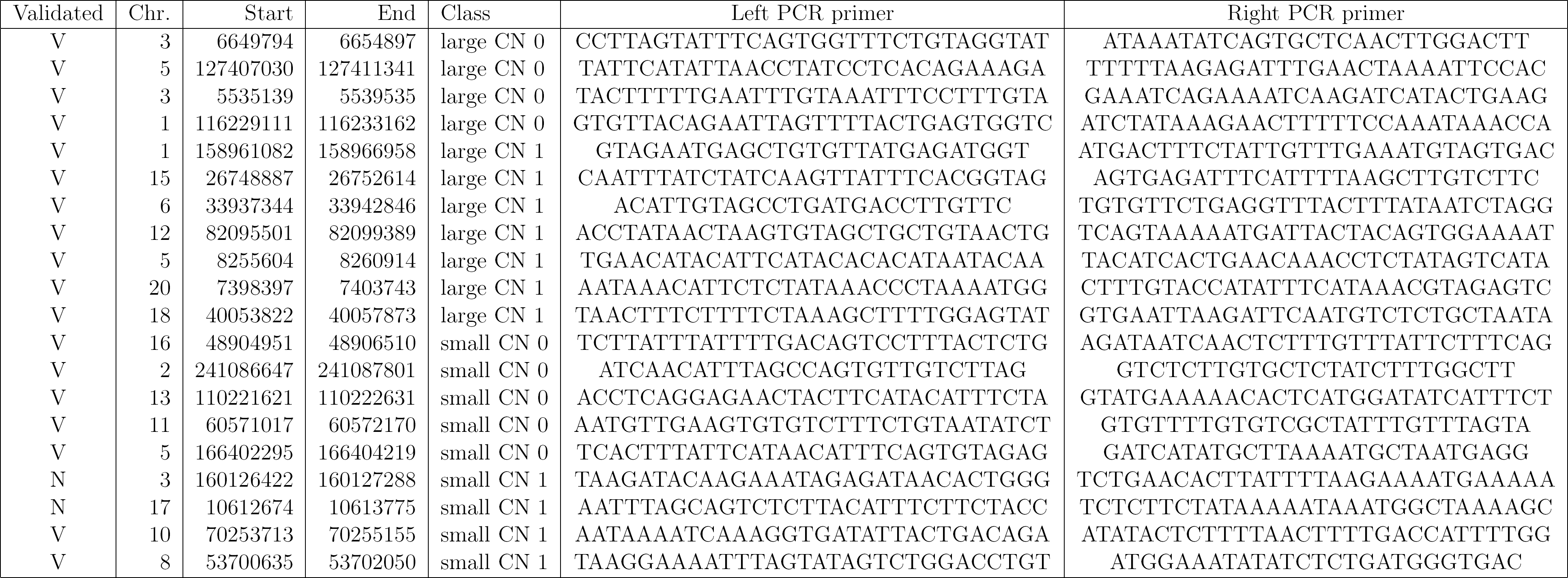
Experimental validation results. Location of the validated (V) and non-validated (N) CNVs for different classes. The last two columns show the primer sequences used for PCR amplification.

**Table S2:**
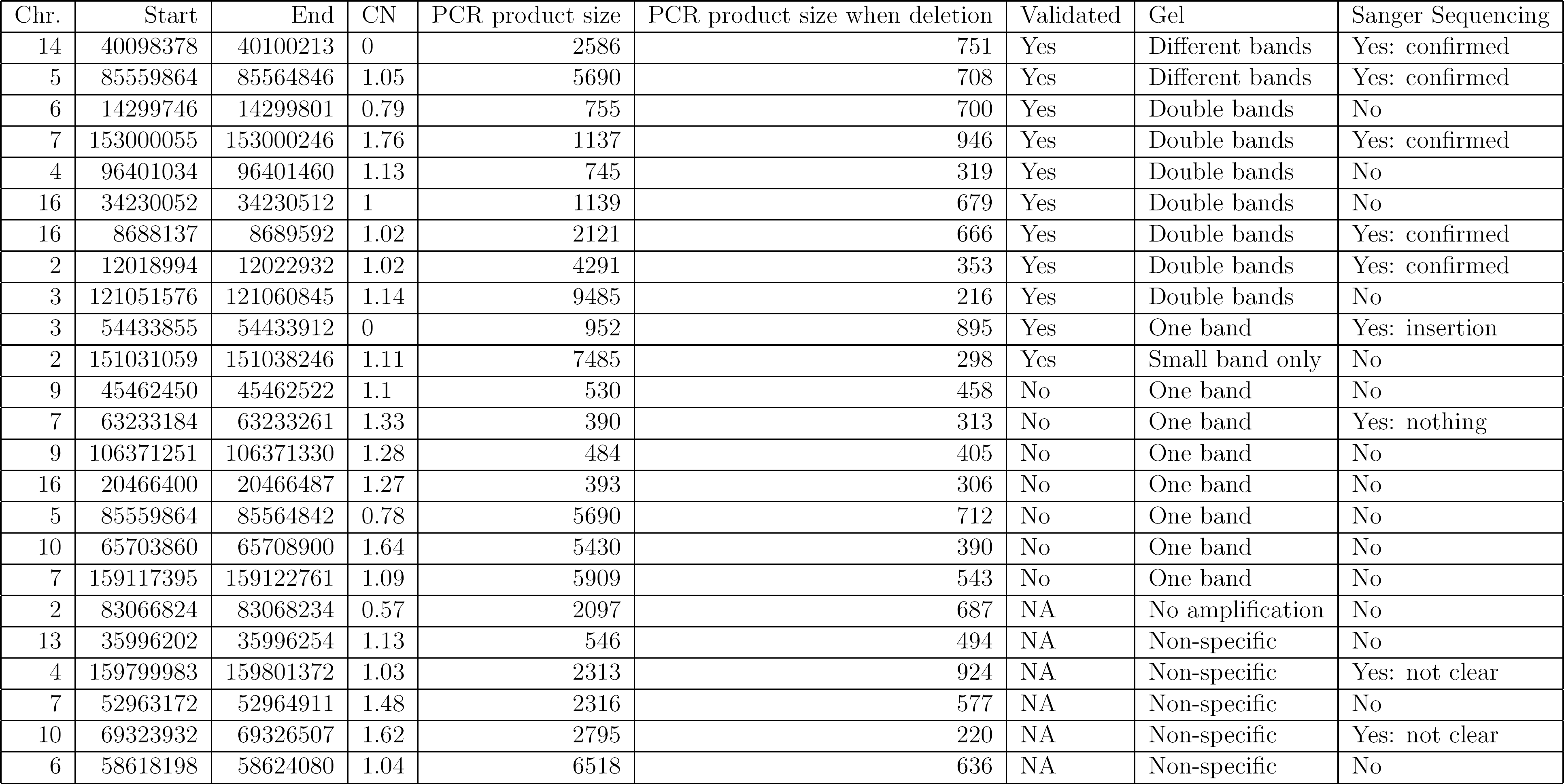
Experimental validation in low-coverage regions. The result of the PCR validation was either concordant with `PopSV` call (Yes), discordant (No) or inconclusive (NA). In some cases, Sanger sequencing was performed. The *CN* column is the estimated copy-number of the deleted allele.

| Chr. | Start | End | CN | PCR product size | PCR product size when deletion | Validated | Gel | Sanger Sequencing |
| --- | --- | --- | --- | --- | --- | --- | --- | --- |
| 14 | 40098378 | 40100213 | 0 | 2586 | 751 | Yes | Different bands | Yes: confirmed |
| 5 | 85559864 | 85564846 | 1.05 | 5690 | 708 | Yes | Different bands | Yes: confirmed |
| 6 | 14299746 | 14299801 | 0.79 | 755 | 700 | Yes | Double bands | No |
| 7 | 153000055 | 153000246 | 1.76 | 1137 | 946 | Yes | Double bands | Yes: confirmed |
| 4 | 96401034 | 96401460 | 1.13 | 745 | 319 | Yes | Double bands | No |
| 16 | 34230052 | 34230512 | 1 | 1139 | 679 | Yes | Double bands | No |
| 16 | 8688137 | 8689592 | 1.02 | 2121 | 666 | Yes | Double bands | Yes: confirmed |
| 2 | 12018994 | 12022932 | 1.02 | 4291 | 353 | Yes | Double bands | Yes: confirmed |
| 3 | 121051576 | 121060845 | 1.14 | 9485 | 216 | Yes | Double bands | No |
| 3 | 54433855 | 54433912 | 0 | 952 | 895 | Yes | One band | Yes: insertion |
| 2 | 151031059 | 151038246 | 1.11 | 7485 | 298 | Yes | Small band only | No |
| 9 | 45462450 | 45462522 | 1.1 | 530 | 458 | No | One band | No |
| 7 | 63233184 | 63233261 | 1.33 | 390 | 313 | No | One band | Yes: nothing |
| 9 | 106371251 | 106371330 | 1.28 | 484 | 405 | No | One band | No |
| 16 | 20466400 | 20466487 | 1.27 | 393 | 306 | No | One band | No |
| 5 | 85559864 | 85564842 | 0.78 | 5690 | 712 | No | One band | No |
| 10 | 65703860 | 65708900 | 1.64 | 5430 | 390 | No | One band | No |
| 7 | 159117395 | 159122761 | 1.09 | 5909 | 543 | No | One band | No |
| 2 | 83066824 | 83068234 | 0.57 | 2097 | 687 | NA | No amplification | No |
| 13 | 35996202 | 35996254 | 1.13 | 546 | 494 | NA | Non-specific | No |
| 4 | 159799983 | 159801372 | 1.03 | 2313 | 924 | NA | Non-specific | Yes: not clear |
| 7 | 52963172 | 52964911 | 1.48 | 2316 | 577 | NA | Non-specific | No |
| 10 | 69323932 | 69326507 | 1.62 | 2795 | 220 | NA | Non-specific | Yes: not clear |
| 6 | 58618198 | 58624080 | 1.04 | 6518 | 636 | NA | Non-specific | No |

**Table S3:**
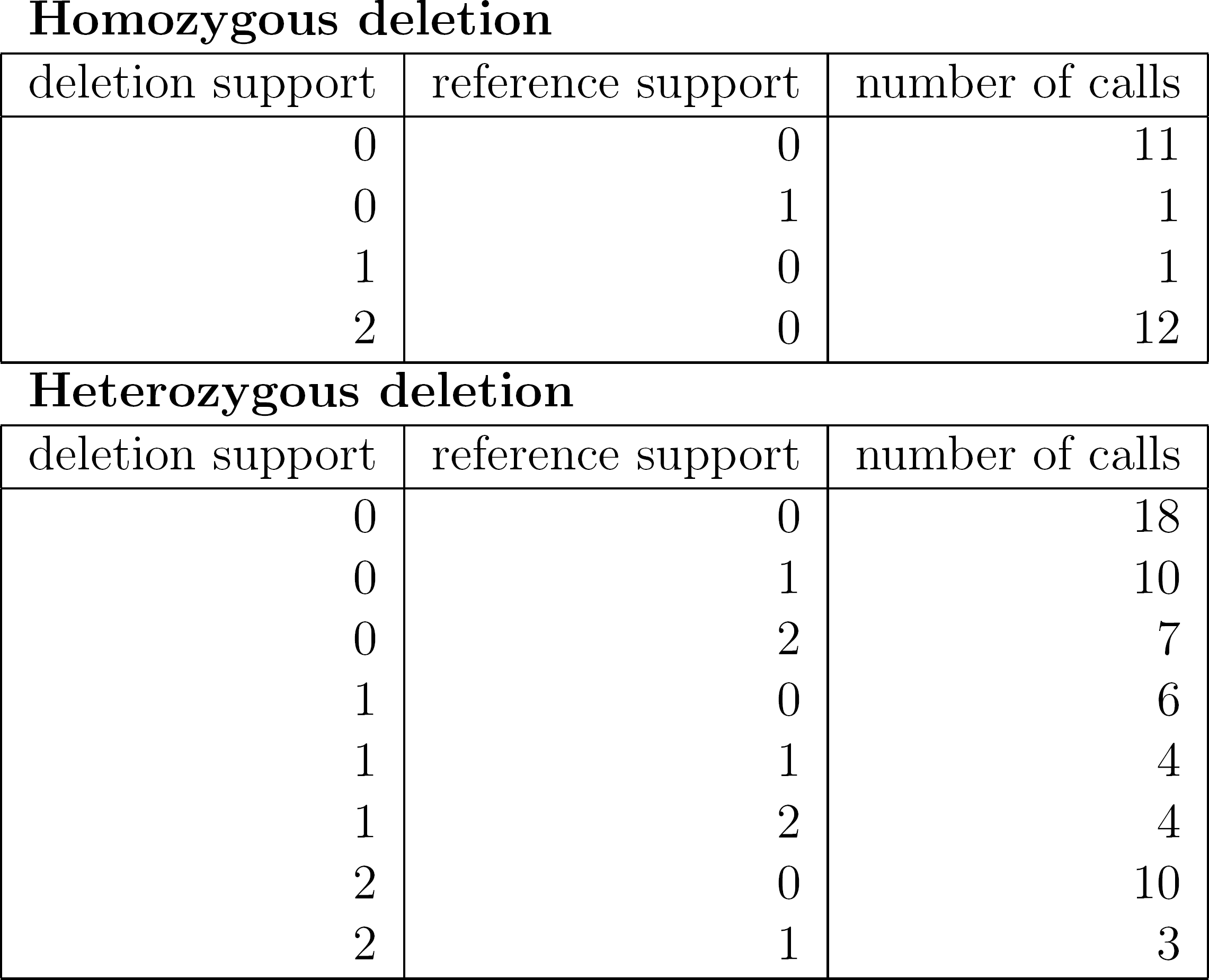
Investigating low-mappability deletion calls with two CEPH12878 assemblies. The first two columns represent the number of assemblies (0, 1 or 2) supporting the deleted allele or the reference allele. The third column shows the number of `PopSV` calls in each category.

Homozygous deletion
| deletion support | reference support | number of calls |
| --- | --- | --- |
| 0 | 0 | 11 |
| 0 | 1 | 1 |
| 1 | 0 | 1 |
| 2 | 0 | 12 |

| deletion support | reference support | number of calls |
| --- | --- | --- |
| 0 | 0 | 18 |
| 0 | 1 | 10 |
| 0 | 2 | 7 |
| 1 | 0 | 6 |
| 1 | 1 | 4 |
| 1 | 2 | 4 |
| 2 | 0 | 10 |
| 2 | 1 | 3 |

**Table S4:** Deletions, duplications and CNVs in the 1000 Genomes Project. We removed variants with high frequency (> 80%), variants in the chromosome X, and variants smaller than 300 bp in order to compare with `PopSV`’s numbers (Table 1). WG: whole genome; ELC: extremely low-coverage regions. The *Total* number of variants is the total number after collapsing recurrent variants. *Affected genome* represents the amount of the reference genome that overlaps at least one CNV.

| Sample | Type | Total variants | Variants per sample |  | Avg Size (Kbp) | Variants < 3 Kbp<br>Per sample | Affected genome (Mbp) |  |
| --- | --- | --- | --- | --- | --- | --- | --- | --- |
|  |  |  | *WG* | *ELC* |  |  | Total | Per sample |
| 2,504 | All | 41,979 | 1,024.44 | 2.22 | 6.00 | 700.52 | 580.03 | 6.14 |
|  | DEL | 36,102 | 975.32 | 2.22 | 4.67 | 700.52 | 342.97 | 4.56 |
|  | DUP | 8,503 | 48.26 | 0.00 | 32.54 | 0.00 | 331.48 | 1.57 |

## 10 SUPPLEMENTARY FIGURES

**Figure S1:**
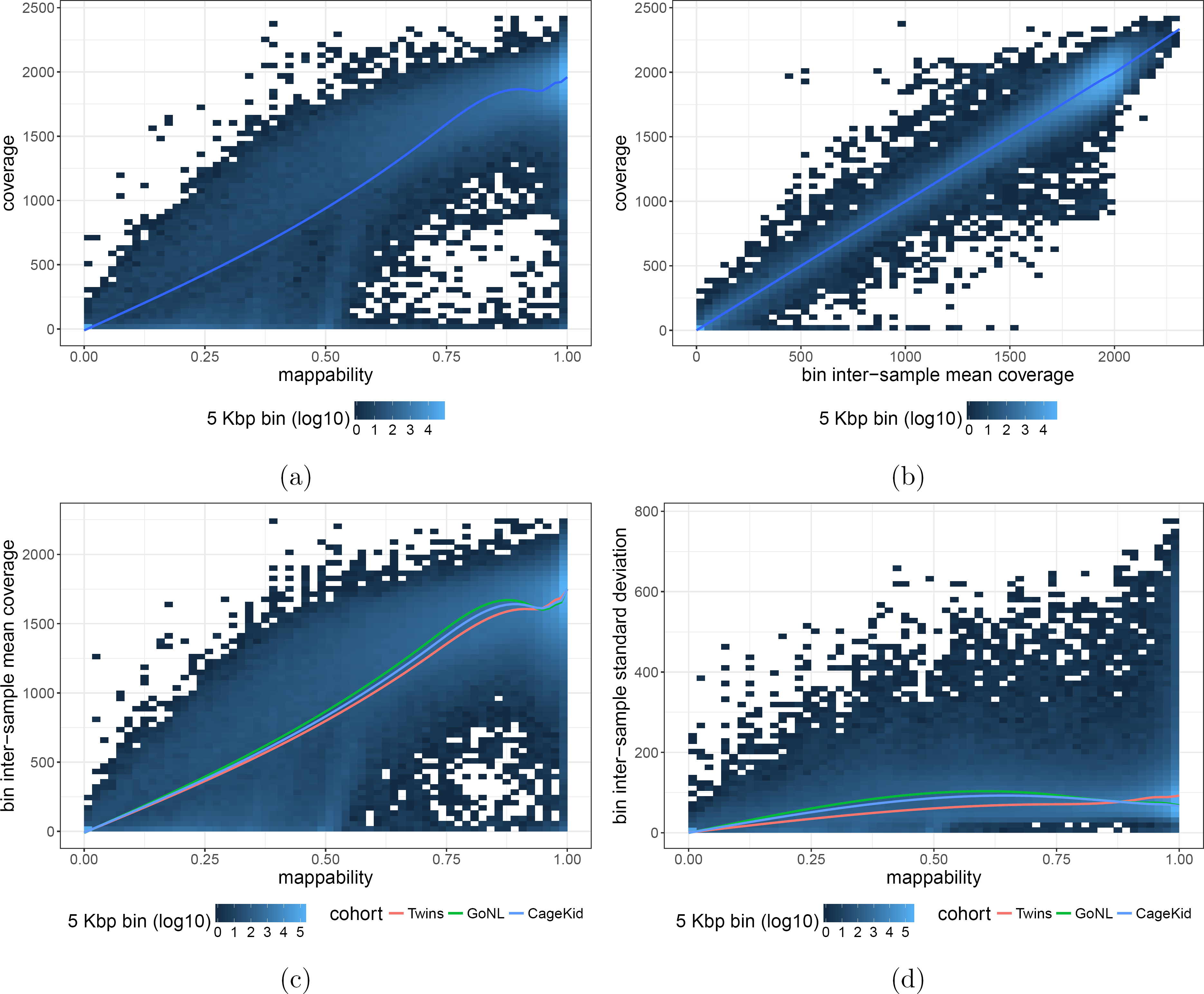
Coverage, mappability and population-based measures. a-b) Read coverage in a sample (y-axis) versus mappability (a) or the inter-sample average coverage (b). c-d) Inter-sample mean (c) and standard deviation (d) were fitted against the mappability in each cohort separately. The tiles represent all cohorts pooled together.

**Figure S2.**
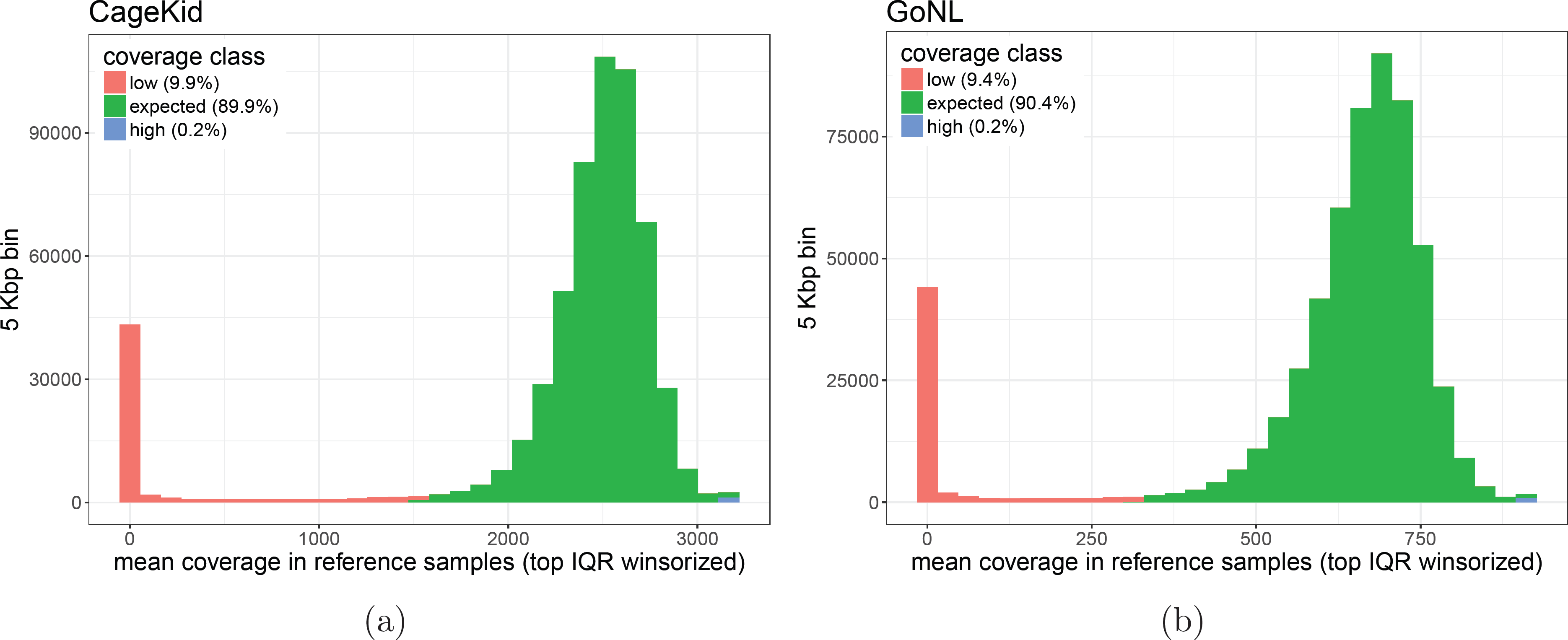
Average coverage in reference samples in the CageKid (a) and GoNL (b) datasets.

**Figure S3:**
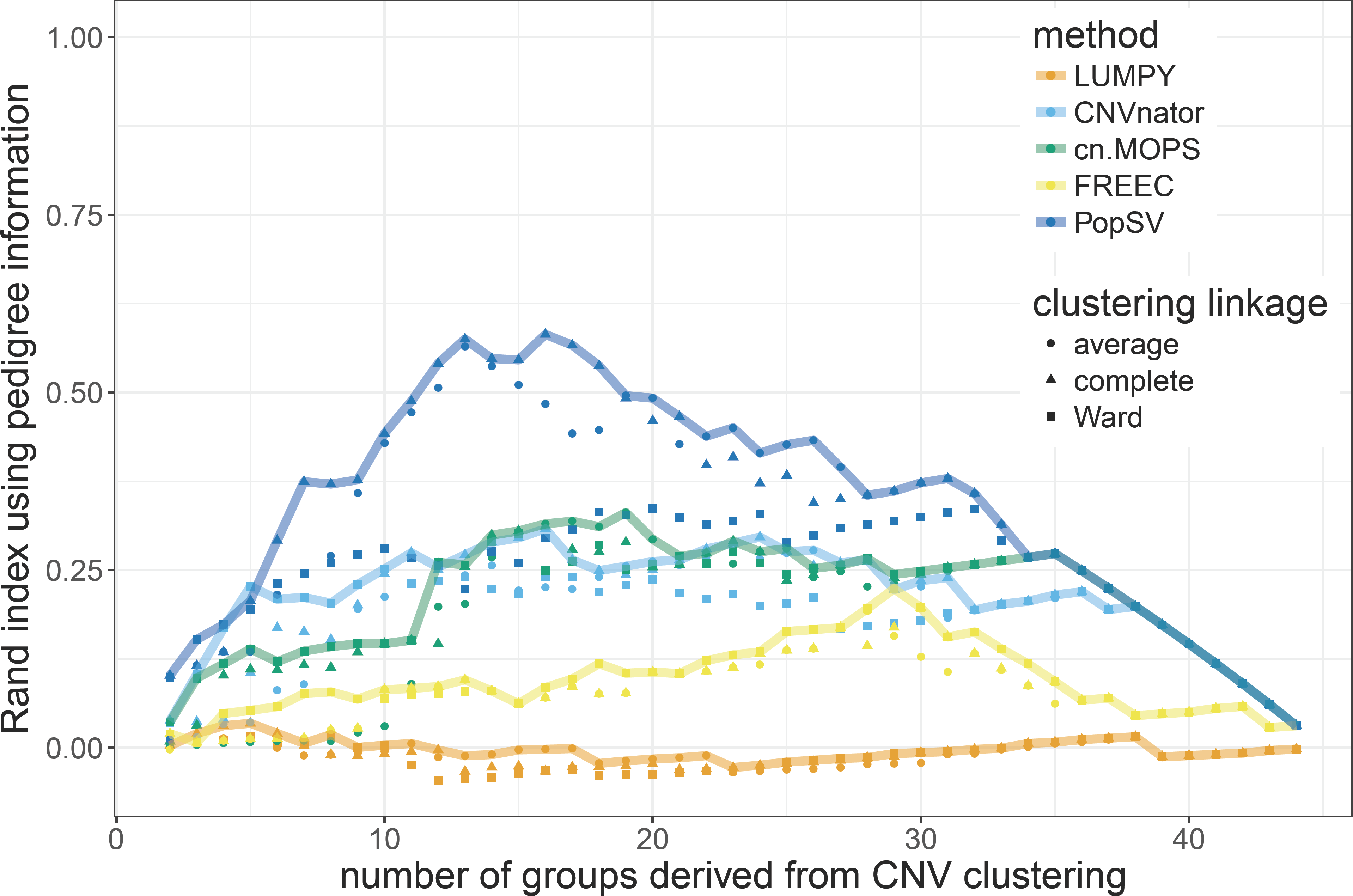
Rand index between the pedigree information and the dendogram from CNV calls in low-coverage regions. The dendogram for CNV-based clustering was cut at different levels (x-axis) and the groups compared to the pedigree (family-level) with the Rand index (y-axis). For each method, the line highlights the best performance across three linkage criteria.

**Figure S4:**
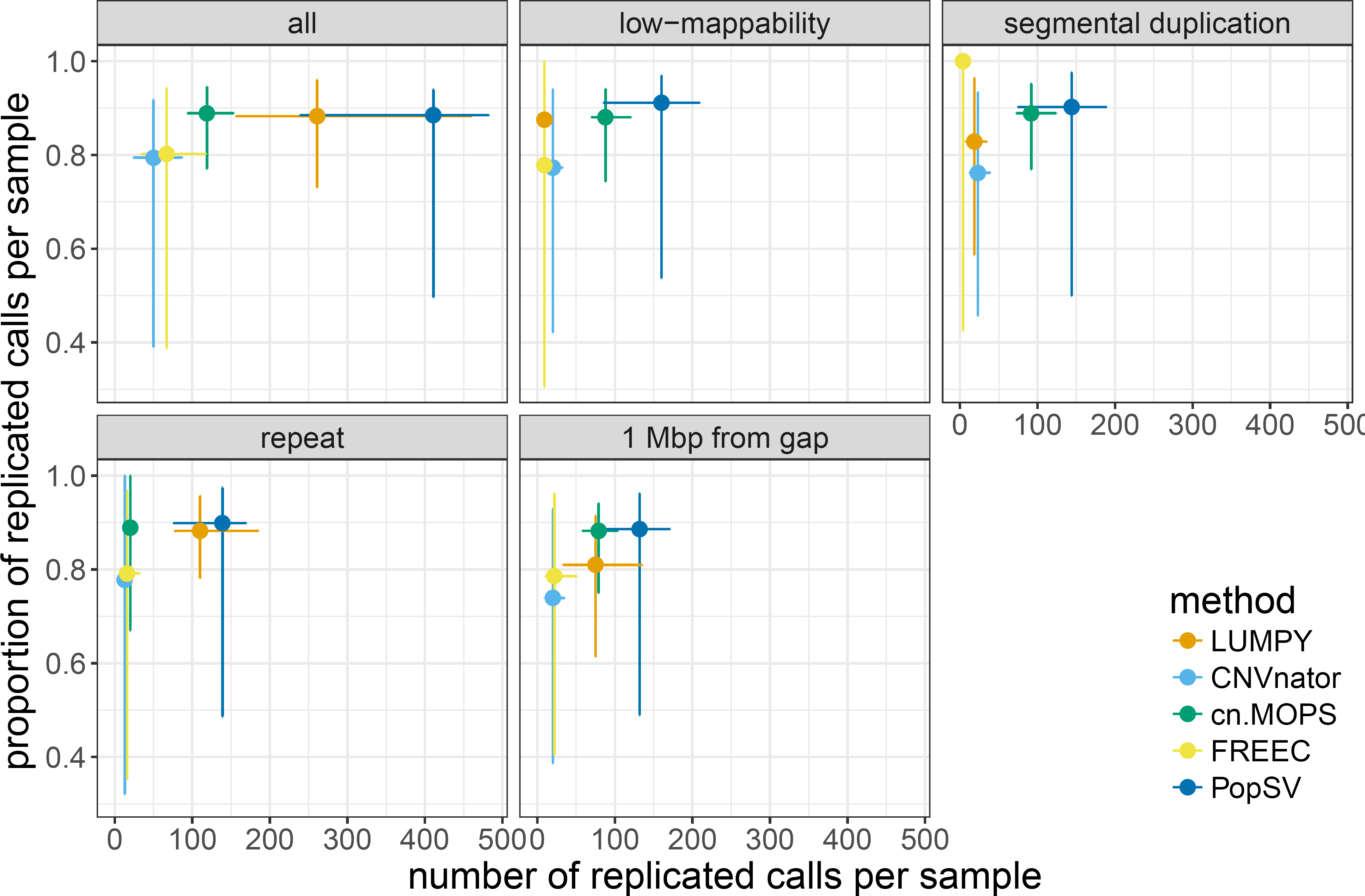
PopSV’s performance in low-mappability regions in CageKid dataset. Proportion and number of calls replicated in the paired tumor. The point shows the median value per sample, the error bars the 95% confidence interval.

**Figure S5:**
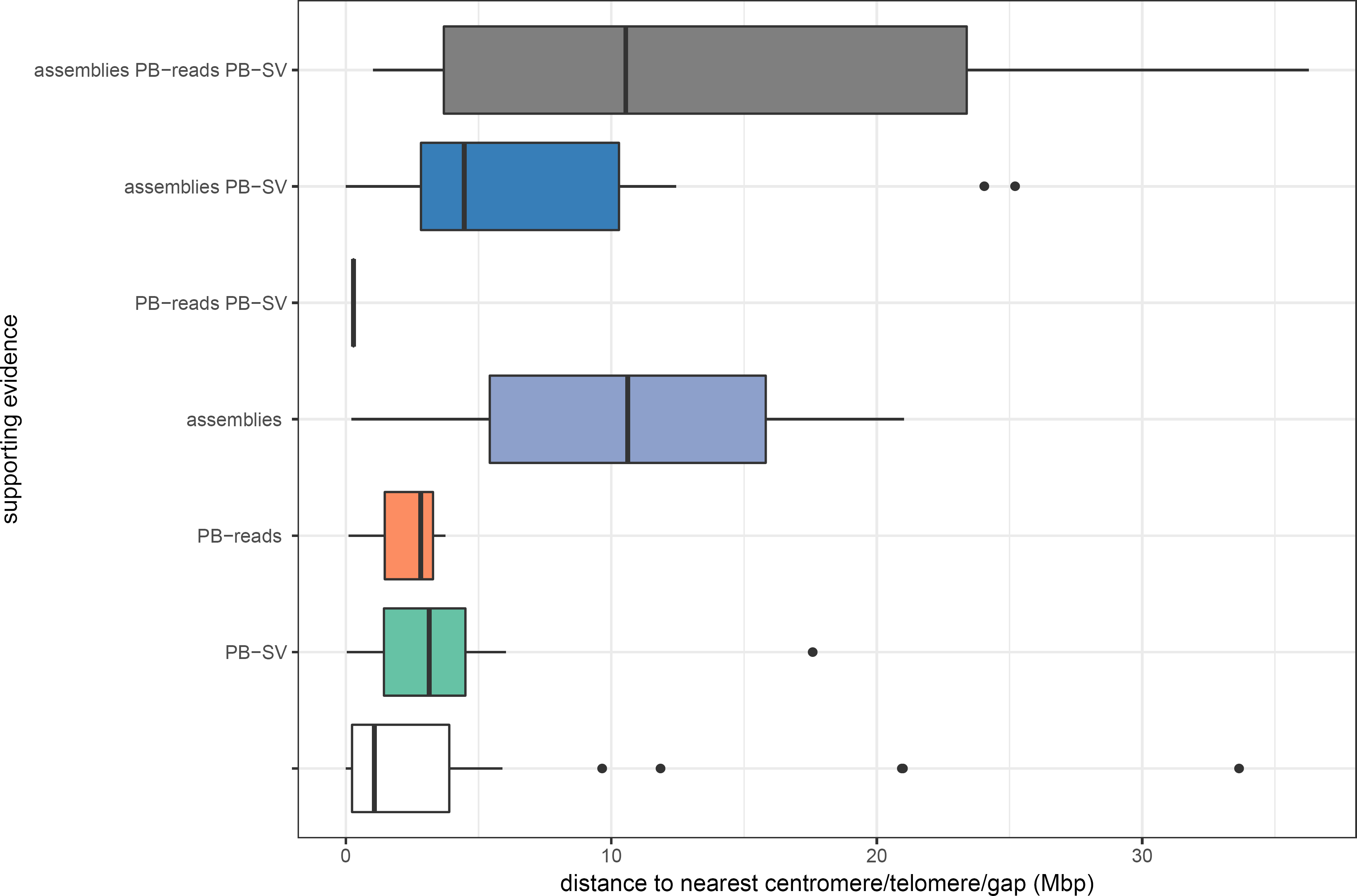
Distance to assembly gaps and supporting evidence from long-read sequencing in CEPH12878. Deletions in low-mappability regions were grouped by their supporting evidence (y-axis and colors). *assemblies*: deletion observed in at least one of the two public assemblies. *PB-SV:* overlap with a structural variant called from the PacBio reads^53^. *PB-reads:* deletion observed in the local assembly or consensus of the PacBio reads. Variants with no support are represented by the white boxplot.

**Figure S6:**
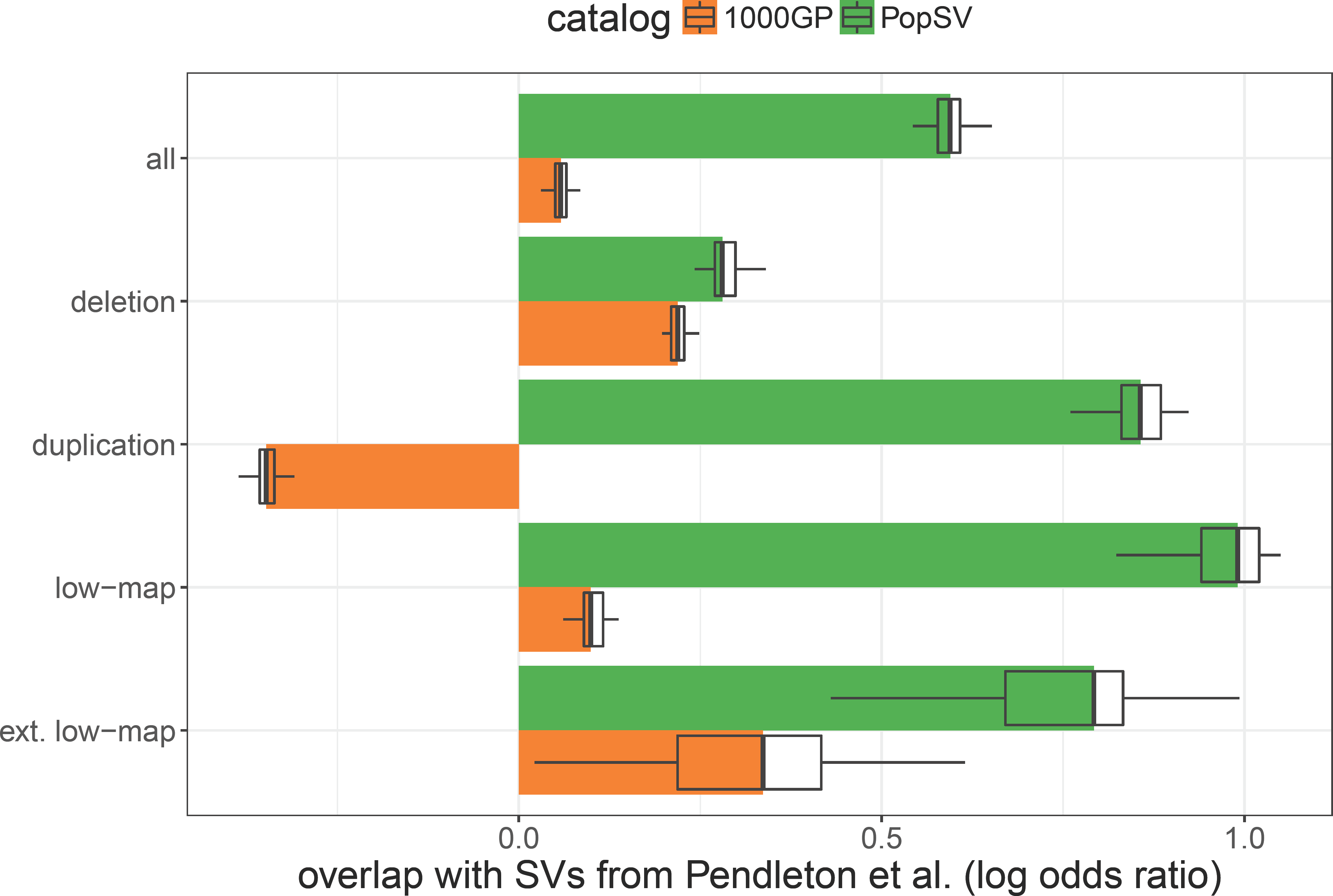
Overlap between PopSV catalog and calls from Pendleton et al. Recurrent calls were collapsed in each catalog (i.e `PopSV` and the 1000 Genomes Project (1000GP)). The proportion of the collapsed calls overlapping calls from Pendleton et al.^53^ was computed. The fold-enrichment is produced by drawing control regions with similar size distribution as Pendleton’s calls. *low-map:* calls in low-mappability regions; *ext. low-map:* calls in extremely low-mappability regions.

**Figure S7:**
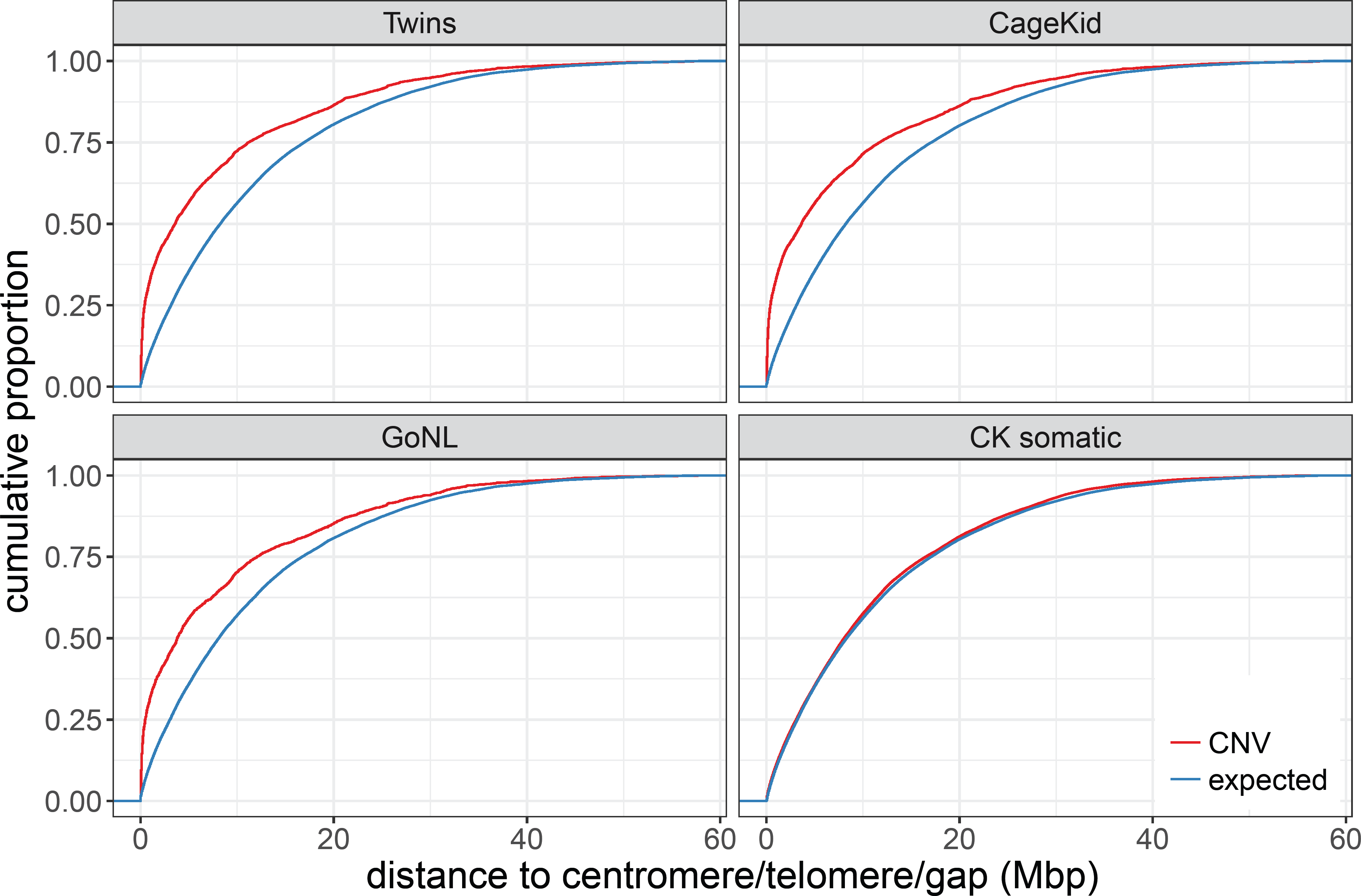
Distance to a centromere, telomere or assembly gap. The y-axis represents the cumulative proportion of the affected genome. The *expected* curve is computed from uniformly distributed genomic regions with matched size.

**Figure S8:**
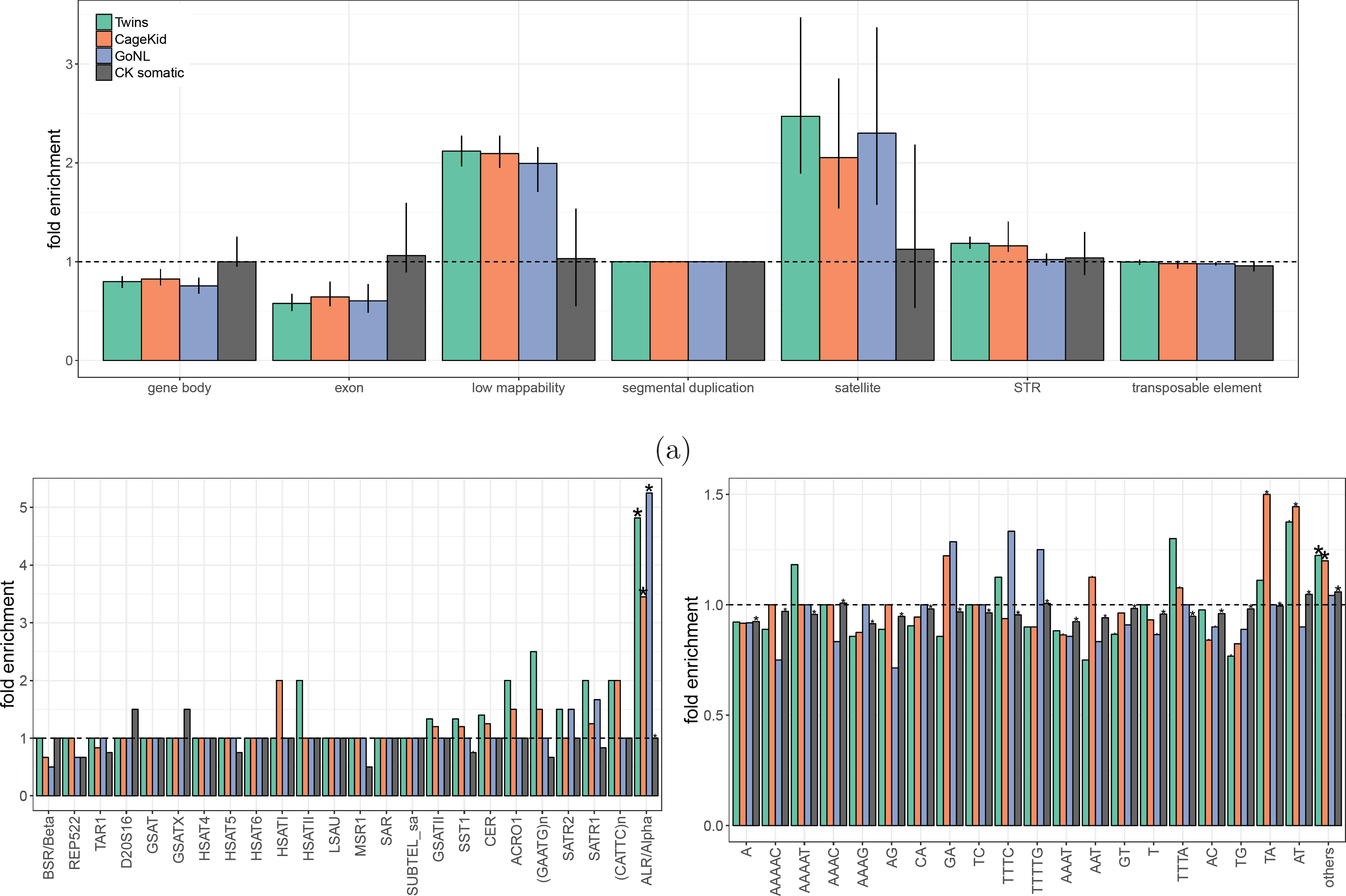
CNVs enrichment after controlling for segmental duplication overlap and distance to CTG. Enrichment of CNVs in a) different genomic features, b) satellite families and c) simple repeats in the different cohorts (colors). Bars show the median fold enrichment across samples compared to control regions. The star represents significant enrichment from the logistic regression.

**Figure S9:**
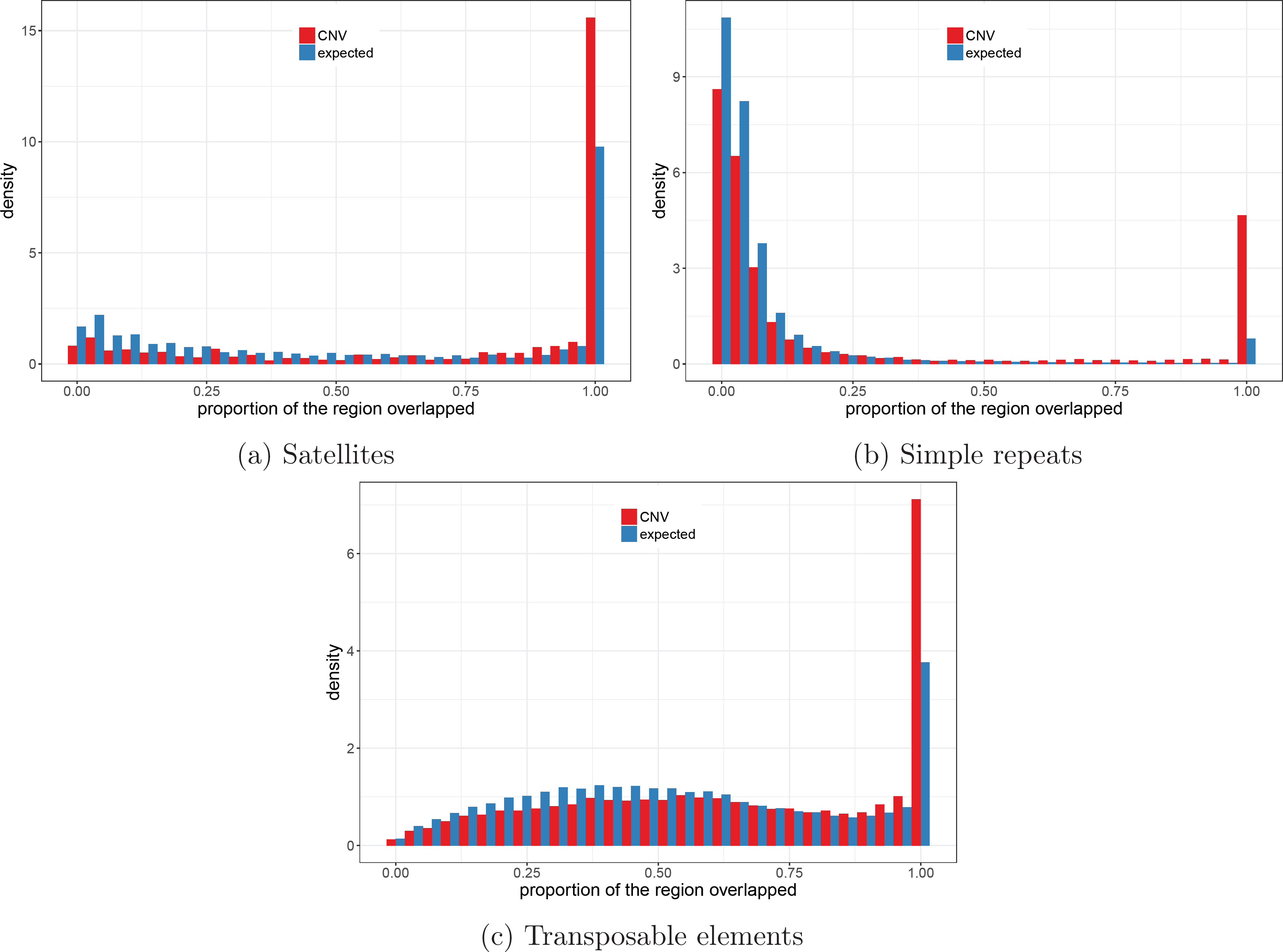
Overlap between CNVs and repeats. The histograms represent the proportion of the CNV region that overlaps a) a satellite, b) a simple repeat or c) a transposable element, when they do overlap. The *expected* distribution is computed from the control regions used for the enrichment analysis.

**Figure S10:**
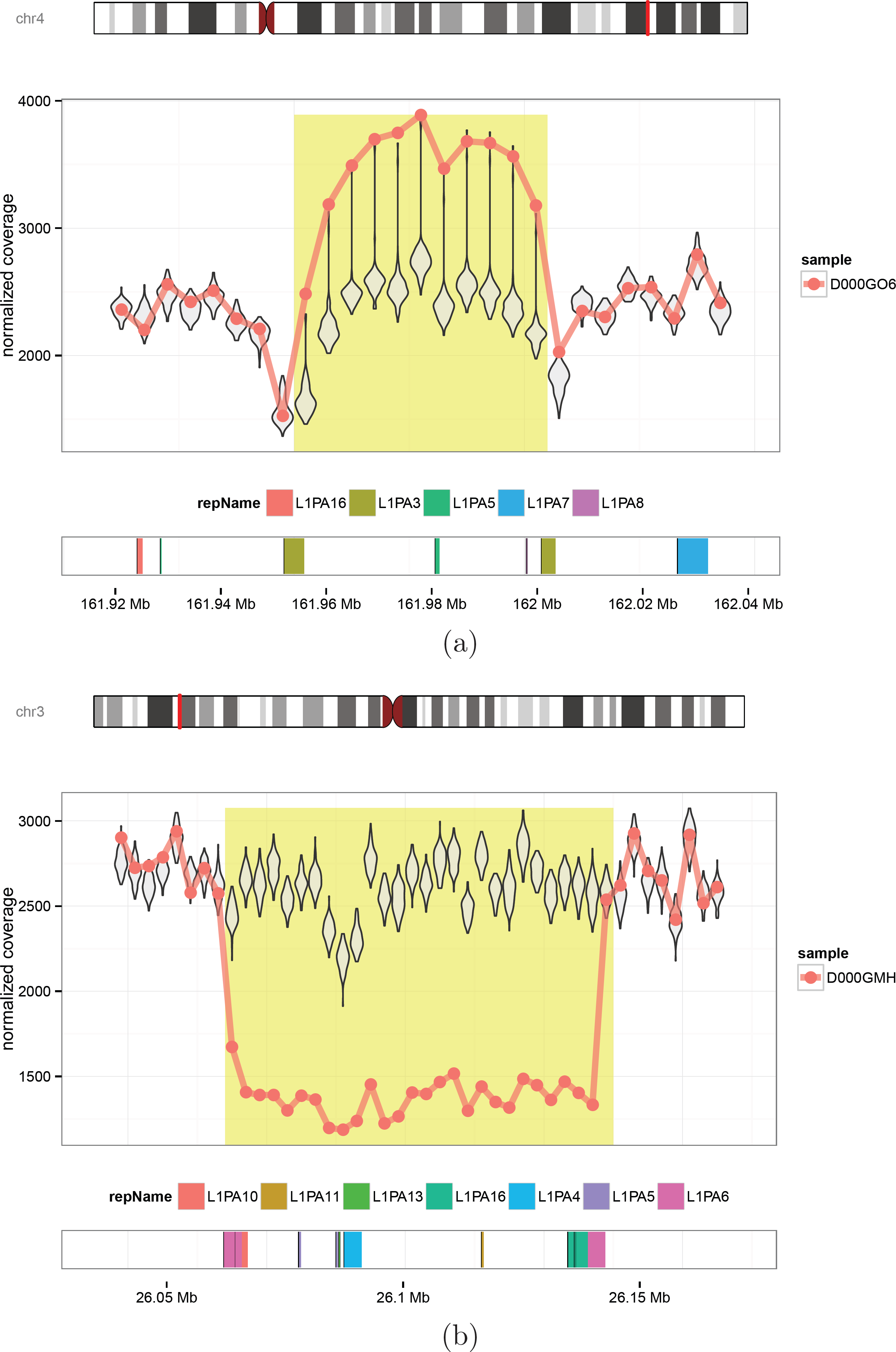
Polymorphism likely caused by non-homologous allelic recombination between L1PA repeats. Examples of CNV likely caused by non-allelic homologous recombination between two L1PA3 repeats (a) or L1PA6 (b). The line and points represent the coverage of one sample with a duplication (a) or a deletion (b), highlighted in yellow; the violin plots represent the distribution of the coverage in the reference samples.

## 11 SUPPLEMENTARY INFORMATION

### 11.1 Data

#### Twin study

All patients gave informed consent in written form to participate in the Quebec Study of Newborn Twins^47^. Ethic boards from the Centre de Recherche du CHUM, from the UniversitÃľ Laval and from the Montreal Neurological Institute approved this study. Sequencing was done on an Illumina HiSeq 2500 (paired-end mode, fragment length 300 bp). The reads were aligned using a modified version of the Burrows-Wheeler Aligner (`bwa` version 0.6.2-r126-tpx with threading enabled). The options were `’bwa aln -t 12 -q 5’` and `’bwa sampe -t 12’`. The aligned reads are available on the European Nucleotide Archive under ENA PRJEB8308. The 45 samples had an average sequencing depth of 40x (minimum 34x / maximum 57x).

#### Renal cell carcinoma

WGS data from renal cell carcinoma is presented in details in the CageKid paper^48^. In short, 95 pairs of normal/tumor tissues were sequenced using GAIIx and HiSeq2000 instruments. Paired-end reads of size 100 bp totaled an average sequencing depth of 54x (minimum 26x / maximum 164x). Reads were trimmed with `FASTX-Toolkit` and mapped per lane with BWA backtrack to the GRCh37 reference genome. `Picard` was used to adjust pairs coordinates, flag duplicates and merged lane. Finally, realignment was done with `GATK`. Raw sequence data have been deposited in the European Genome-phenome Archive, under the accession code EGAS00001000083.

#### Genome of the Netherlands

WGS data from the GoNL project is described in details in Francioli et al.^34^. This data have been derived from different sample collections:

- The LifeLines Cohort Study, supported by the Netherlands Organization of Scientific Research (NWO, grant 175.010.2007.006), the Dutch governmentâĂŹs Economic Structure Enhancing Fund (FES), the Ministry of Economic Affairs, the Ministry of Education, Culture and Science, the Ministry for Health, Welfare and Sports, the Northern Netherlands Collaboration of Provinces (SNN), the Province of Groningen, the University Medical Center Groningen, the University of Groningen, the Dutch Kidney Foundation and Dutch Diabetes Research Foundation.
- The EMC Ergo Study.
- The LUMC Longevity Study, supported by the Innovation-Oriented Research Program on Genomics (SenterNovem IGE01014 and IGE05007), the Centre for Medical Systems Biology and the National Institute for Healthy Ageing (Grant 05040202 and 05060810).
- VU Netherlands Twin Register. In short, samples were sequenced on an Illumina HiSeq 2000 instrument (91-bp paired-end reads, 500-bp insert size). We downloaded the aligned read sequences (BAM) for the 500 parents in the data set. We further performed indel realignment using GATK 3.2.2, adjusted pairs coordinates with `Samtools` 0.1.19, marked duplicates with `Picard` 1.118, and performed base recalibration (`GATK` 3.2.2). The average sequencing depth was 14x (minimum 9x / maximum 59x).

#### Genomic annotations

Gencode annotation (V19) was directly downloaded from the consortium FTP server at ftp://ftp.sanger.ac.uk/pub/gencode/Gencode_human/release_19/gencode.v19.annotation.gtf.gz. Other genomic annotations were downloaded from the UCSC database^58^ server at http://hgdownload.soe.ucsc.edu/goldenPath/hg19/database. The file names of the corresponding annotations are

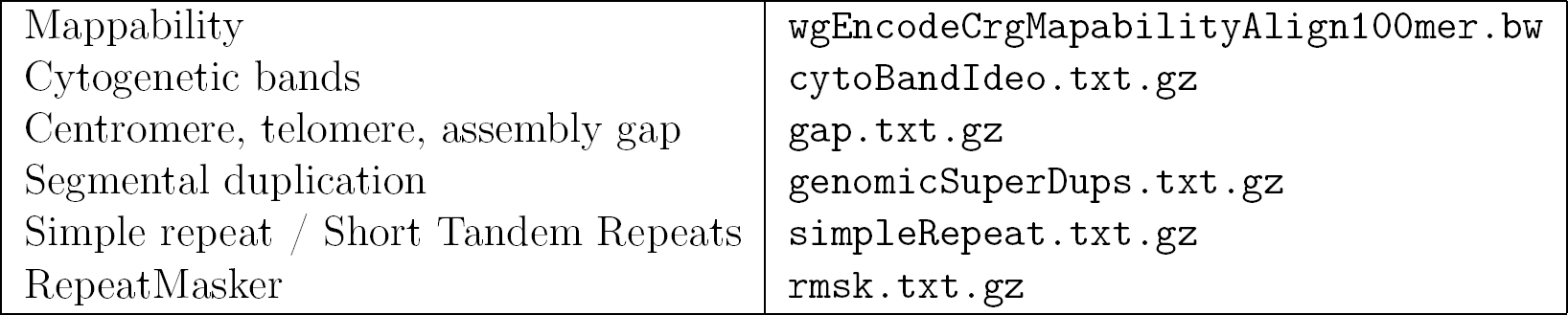

|  |  |
| --- | --- |
| Mappability | `wgEncodeCrgMapabilityAlign100mer.bw` |
| Cytogenetic bands | `cytoBandIdeo.txt.gz` |
| Centromere, telomere, assembly gap | `gap.txt.gz` |
| Segmental duplication | `genomicSuperDups.txt.gz` |
| Simple repeat / Short Tandem Repeats | `simpleRepeat.txt.gz` |
| RepeatMasker | `rmsk.txt.gz` |

### 11.2 Read count across the genome

The genome was fragmented in non-overlapping bins of fixed size. The number of properly mapped reads was used as a coverage measure, defined as read pairs with correct orientation and insert size, and a mapping quality of 30 (Phred score) or more. In each sample, GC bias was corrected by fitting a LOESS model between the bin’s coverage and the bin’s GC content. For each bin, the correction factor was computed as the mean coverage across all the bins divided by the predicted coverage from the LOESS model and the GC content of the bin. We used a bin size of 5 Kbp for most of the analysis. When specified, we used a smaller bin size of 500 bp.

### 11.3 RD and mappability estimates

To investigate the bias in RD we used the read counts in 5 Kbp bins. Bins with extremely high coverage were identified and removed when deviating from the median coverage by more than 5 standard deviation. First the coverage of the 45 samples from the Twin study were combined and quantile normalized. At that point the different samples had the same global coverage distribution and no bins with extreme coverage or GC bias.

The mappability track^27^ was downloaded from UCSC^58^ (`wgEncodeCrgMapabilityAlign100mer.bw`) and the average mappability was computed for each bin. One sample was randomly selected and we compared its coverage with the mappability estimates. We then computed the mean and standard deviation of the coverage in each bin across the other samples and compared it with the sample coverage. We also compared the inter-sample average with the mappability estimates.

To compute Z-scores that integrates the observed coverage variation we used two approaches. The first modeled the coverage metrics (average or standard deviation) using the mappability estimates and computed a Z-score from the predicted coverage and global standard deviation. A generalized additive model was fitted using a cubic regression spline on the mappability estimates (`mgcv` R package). In the second approach, Z-scores were computed using the inter-sample average and standard deviation. The normality of these two Z-score distributions were compared in term of excess kurtosis and skewness. For the kurtosis and skewness computation, we removed outlier Z-scores with an absolute value greater than 10. These bins could be regions of CNV and would bias the estimates. The Z-score distributions were also compared in bins from 10 different mappability intervals.

We repeated this analysis pooling 45 samples from each of the three datasets. After quantile normalization, the inter-sample coverage mean and standard deviation were computed separately in each cohort and compared with the mappability estimates.

### 11.4 CNV detection with PopSV

#### Binning the genome

We ran two separate analysis on the three datasets. Bin sizes of 5 Kbp and 500 bp were used on the Twin study and renal cell carcinoma. Because of its lower sequencing depth, the 500 bp run on GoNL gave only partial results. More precisely, we observed a truncated distribution of the copy-number estimates, with most of the 1 and 3 copy number variants missing. It means that at this resolution many one-copy variation cannot be differentiated from background noise. For this reason we ran GoNL analysis using 2 Kbp and 5 Kbp bins.

#### Constructing the set of reference samples

In each dataset we choose the reference samples as follows: in the renal cancer dataset from the normal samples, in the Twin study from all the samples, in GoNL from a subset of 200 samples (see below). For each dataset, a Principal Component Analysis (PCA) was performed across samples on the counts normalized globally (median/variance adjusted). The resulting first two principal components are used to verify the homogeneity of the reference samples. Although our three datasets showed different levels of homogeneity, we didn’t need to exclude samples or split the analysis. The effect of weak outlier samples was either corrected by the normalization step or integrated in the population-view.

In GoNL, we decided to use only 200 of the 500 samples as reference. They were selected to span a maximum of the space defined by the principal components. In contrast to random selection, this ensures that weak outliers are included in the final set of reference samples, hence maximizing the technical variation integrated in the population-view.

Moreover, the principal components were used to select one control sample from the final set of reference samples. This sample is used in the normalization step as a baseline to normalize other samples against. We picked the sample closest to the centroid of the reference samples in the Principal Component space.

#### CNV calling

After targeted normalization the coverage in each sample is compared to the coverage in the reference samples. A Z-score is computed and translated into a P-value that is then corrected for multiple testing. Consecutive bins with significant excess or lack of reads are merged and returned as potential duplication or deletion. Copy number estimates are derived from the coverage across the bin and the average coverage across the reference samples. However, it is important to note that the definition of a variant is different from other methods. Here a variant is defined by the major allele in the population rather than the reference genome state. Most of the genome is in a diploid state compared to the reference genome and sufficiently covered by sequencing reads that the copy number state can be correctly estimated by `PopSV`’s population-based approach. However, highly polymorphic variants are called relative to the major allele in the population and additional efforts are required to assess the copy number state. Variants in extremely low-mappability regions are also difficult to fully characterize and might be caused by rare insertion in the reference genome or complex alleles. Nonetheless, `PopSV` can efficiently detect the presence of CNV in any situation. More details are available in the method paper (Monlong et al., under review).

#### Coverage tracks

For each run, we constructed coverage tracks based on the average coverage in the reference samples. Bins where the reference samples had, on average, the expected coverage were classified as *expected coverage*. Bins with a coverage lower than 4 standard deviation from the median were classified as *low-mappability(or low coverage).* To ensure robustness, the standard deviation was derived from the Median Absolute Deviation. We use regions with low coverage to define *low-mappability regions*, as the low coverage is a result of the lower mappability of a region. Because the standard deviation is used, the number of regions classified as *low-mappability* is lower in datasets with more RD variance.

Eventually, we also defined *extremely low coverage* region which have an average coverage below 100. This sub-class of *low coverage* region was used in a few analyses to highlight the most challenging regions.

Regions were annotated with the overlap with protein-coding genes and segmental duplications (see Genomic annotations), and the distance to the nearest centromere, telomere or assembly gap. Finally, we computed the number of protein-coding genes overlapping at least one low-coverage region.

### 11.5 Validation and benchmark

#### Running

`FREEC`, `CNVnator`, `cn.MOPS` and `LUMPY FREEC`^16^ segments the RD values of a sample using a LASSO-based algorithm. It was run on each sample separately, starting from the BAM file, using the same bin sizes as for `PopSV`. `FREEC` internally corrects the RD for GC and mappability bias. In order to compare its performance in low-mappability region, the minimum *“telocentromeric”* distance was set to 0. The remaining parameters were set to default. Of note an additional run with slightly looser parameter (`breakPointThreshold=0.6`) was performed to get a larger set of calls used in some parts of the *in silico* validation analysis to deal with borderline significant calls.

`CNVnator`^17^ uses a mean-shift technique inspired from image processing. It was run on each sample separately, starting from the BAM file, using the same bin sizes as for `PopSV`. `CNVnator` also corrects internally for GC bias and we used default parameters. For the analysis using higher confidence calls, we used calls with either ‘eval1’ or ‘eval2’ lower than 10^-5^ (instead of the default 0.05).

`cn.MOPS`^18^ considers simultaneously several samples and detects copy number variation using a Poisson model and a Bayesian approach. It was run on the same GC-corrected bin counts used for `PopSV`. All the samples are analyzed jointly. Of note an additional run with slightly looser parameter (`upperThreshold=0.32 and lowerThreshold=-0.42`) was performed to get a larger set of calls used in some parts of the *in silico* validation analysis to deal with borderline significant calls.

`LUMPY`^52^ which uses an orthogonal mapping signal: the insert size, orientation and split mapping of paired reads. The discordant reads were extracted from the BAMs using the recommended commands. Split-reads were obtained by running `YAHA`^51^ with default parameters. All the CNVs (deletions and duplications) larger than 300 bp were kept for the upcoming analysis. Calls with 5 or more supporting reads were considered high-confidence.

#### Clustering samples from the Twin study

A distance between two samples A and B was defined as: 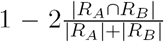 where R_A_ represents the regions called in sample A, 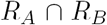 the regions called in both A and B, and |*R*| the cumulative size of the regions. Hence, the similarity between two samples is represented by the amount of sequence found in both divided by the average amount of sequence called. This distance is used for hierarchical clustering of the samples in the Twin dataset. The clustering was performed using only calls in regions with extremely low coverage (reference average ≤100 reads). Different linkage criteria (*average*, *complete* and *Ward*) were used for the exploration. In our dendograms we used the *average* linkage criterion. The concordance between the clustering and the pedigree was estimated by the Rand index, grouping the samples per family. For each method and linkage criteria, the Rand index was computed for every possible dendogram cut (x-axis in Figure S3).

### 11.6 Experimental validation

Experimental validation was performed on samples from the Twin study. In a first validation batch, variants were randomly selected among both one-copy and two-copy deletions. We selected both small (~ 700 bp) and large (~ 4 Kbp) variants in each class. The coverage at base pair resolution was visually inspected for each deletion and, when possible, the breakpoints were fine-tuned. PCR primers were designed to target the whole deleted region. We randomly selected 20 variants out of the variants for which we managed to design PCR primers. We then performed long-range PCR followed by gel electrophoresis. PCR was performed using 50 ng of DNA and the Phusion High-Fidelity DNA Polymerase from Thermo Fisher Scientific: 95 °C 5 minutes followed by 35 cycles (95 °C 30 seconds, 64 °C 30 seconds, 72 °C 45 seconds) and 72 °C 10 minutes. Either a 1% or 1.8% aragose gel was used, depending on the expected size of the amplified fragments. We used a 1 Kb Plus DNA Ladder from Thermo Fisher Scientific.

The presence of a deletion was tested by comparing the size of the amplified fragment in affected and control samples. If the affected sample showed a lower band than a control with a predicted 1 copies, the deletion was considered validated. On the other hand if affected sample and controls had one similar band, the deletion was considered non-validated. Of note, the validation rate might be under-estimated because visual prediction of the breakpoint is not always accurate.

We then randomly selected deletions overlapping low-mappability regions and detected in 6 samples or fewer. We chose to test rare variants because they are likely enriched in false-positives. Hence, this batch of validation represents the most challenging regions to call and validate, and enriched in false-positives. Here we couldn’t use the base-pair coverage to fine-tune the breakpoints because the low-mappability blurs any clear signal. Instead, we retrieved the reads (and their pairs) mapping to the region and assembled them. With this approach we could sometimes get a better breakpoint resolution and design PCR primers that would amplify the deleted region. In addition to gel electrophoresis, the amplified DNA of some regions was sequenced using Sanger sequencing. We randomly selected 17 variants out of the variants for which we managed to design PCR primers.

1 cycle Ãă 95C: 5 minutes 35 cycles: 95C 30 sec, 64C 30 sec, 72C 45 sec 1 cycle: 72C 10 minutes fin Ãă 4C

J’ai essayÃľ des fois le mÃłme protocole mais au lieu de 64C je mettais 65.5C

Le gel est un gel d’agarose 1

L’Ãľchelle c’est ľÃľchelle 1 kb plus ladder de thermofisher.

### 11.7 Analysis of CEPH12878

#### Whole-Genome Sequencing data

High coverage PCR-free Illumina WGS data for 30 samples, including CEPH12878, was downloaded from the 1000 Genomes Project^33^. The ENA accession number is PRJNA260854. The files are also available on the FTP server at http://ftp.1000genomes.ebi.ac.uk/vol1/ftp/release/20130502/supporting/high_coverage_alignments/20141118_high_coverage.alignment.index. Although the sequencing depth is similar to the other datasets (average ~53X), the reads are 250 bp long so the average number of reads per region is lower. Because of the lower read coverage and sample size the CNV calls will be of slightly lower quality. Nonetheless, `PopSV` was run using 5 Kbp bins and all the samples as reference. Using the same coverage track as before we then selected all deletions in CEPH12878 and overlapping low-mappability regions (at least 90% of the call). We then looked for support in public assemblies, SV catalogs and reads from long-read sequencing technologies.

#### Comparison with assemblies

We downloaded the genome assembly produced from short reads, Pacbio and BioNano reads^53^ from ftp://ftp.ncbi.nlm.nih.gov/genomes/all/GCA/001/013/985/GCA_001013985.1_ASM101398v1/GCA_001013985.1_ASM101398v1_genomic.fna.gz. We also downloaded a second assembly that was used 10X Genomics linked reads instead of the Pacbio reads ^54^. It is available at http://kwoklab.ucsf.edu/resources/nmeth_201604_NA12878_hybrid_assembly.fasta.gz.

For each selected variant, we retrieved the two 50 Kbp flanking sequences in the reference genome and aligned them against the public assemblies with BLAST^55^. The output was parsed to identify regions with two flanks aligning in at least 1 Kbp of a contig. MUMmer plots^56^ between the reference sequence and the contigs were visually inspected. The assembly supported `PopSV` calls when a deletion was visible in the expected region (between the flanks). The assembly supported the reference genome sequence when a contig crosses the variant without clear structural variant.

#### SV calls from a long-read sequencing study

We downloaded the SV calls from the Pacbio reads and assembled contigs in Pendleton et al.^53^. The VCF file is publicly available at ftp://ftp-trace.ncbi.nlm.nih.gov/giab/ftp/data/NA12878/NA12878_PacBio_MtSinai/NA12878.sorted.vcf.gz. We overlapped `PopSV` calls with deletions from this SV catalogs. Because we used 5 Kbp bins for `PopSV`, at least 1 Kbp of a `PopSV` calls needed to overlap a deletion from Pendleton et al.^53^ to be considered as sufficient support. Of note, the distribution of the overlap tended to be either null or higher than 1 Kbp supporting this choice.

#### Local assembly of Pacbio reads

Corrected Pacbio reads from Pendleton et al.^53^ were downloaded from ftp://ftp-trace.ncbi.nlm.nih.gov/giab/ftp/data/NA12878/NA12878_PacBio_MtSinai/corrected_reads_gt4kb.fasta. Each read was split in 200 bp fragments and mapped to the human reference genome (version hg19). From this mapping information we selected full Pacbio reads with at least one 200 bp mapping within a region of interest (with 30 Kbp flanks). For each region, the reads were mapped to the reference sequence with exonerate and we kept reads with partial mapping as they may support a SV. These reads were then assembled using `Canu`^67^. A consensus sequence was also derived for reads clustered by alignment breakpoint and the `clustalo`^68^ software. The assembled contigs and consensus were mapped to the reference genome to identify a potential breakpoint. The two regions flanking the alignment breakpoint and the sequence spanning the breakpoint were mapped to the entire genome. We used the results of this genome-wide mapping to select the best candidates: assembled sequence whose flanks align uniquely to the region of interest and with reduced alignment quality for the “middle” sequence that spanned the breakpoint. Candidate contig/consensus were further visualized with MUMmer plots^56^. The assembly supported `PopSV` calls when a deletion was visible in the expected region (between the flanks).

### 11.8 Genomic patterns of CNVs

#### Merging calls from two different bin sizes

Small bins gives better resolution for smaller variants. Large bins gives better sensitivity. For this reason we merged the calls from the 500 bp bin and 5 Kbp bin runs. Variant supported by both sets of calls were merged into one. To decide which set to use for the breakpoints and other information (e.g. copy number estimate), the proportion of overlap was used. If call(s) using small bins overlapped more than a third of a call from the large bin run, it was considered fully recovered by the small bin call which was then used to define breakpoints and other information. If not, the large bin run was considered more appropriate to define the final breakpoints and additional information. Calls unique to each run were simply added to the final set of calls. For the Twin dataset and the renal cancer dataset, calls from the 500 bp and 5Kbp runs were merged. For the GoNL dataset, calls from the 2 Kbp and 5Kbp runs were merged.

#### Computing global estimates of copy number variation

In Table 1, a call in extremely low coverage region is overlapped at more than 90% by the *extremely low coverage* track. To compute the total number of calls, we collapsed calls with an overlap higher than 50%. The amount of sequence affected in a genome was computed by merging all the variants in the cohort and counting the number of affected bases in this reference genome. After the merging step, each base of the genome either overlapped a merged variant or not. Each affected base was counted only once, even if it overlapped CNVs in several samples or with large copy number differences.

#### Comparison with the 1000 Genomes Project SV catalog

The SV catalog from Sudmant et al.^33^ was downloaded from http://ftp.1000genomes.ebi.ac.uk/vol1/ftp/phase3/integrated_sv_map/ALL.wgs.mergedSV.v8.20130502.svs.genotypes.vcf.gz. We retrieved the set of autosomal deletion, duplication and CNVs. When comparing the global estimates of CNV with `PopSV`, we removed deletions smaller than 300 bp as well as variants with high frequency (> 80%). This remaining SVs represent CNVs that could in theory be detected by `PopSV`’s approach. Using this sub-set, we derived the number of variants, number of variants smaller than 3 Kbp, number of variants in *extremely low coverage* regions, and amount of genome affected. These number are computed exactly as the one presented in Table 1 for `PopSV`’s results.

#### CNV frequency comparison

The frequency at which a region is affected by a CNV is computed using calls from the 620 unrelated samples. The copy-number change is not taken into account in the computation and the frequency is derived for all the nucleotide that overlaps at least one CNV. Using each catalog we computed, for each base in the genome, the proportion of individuals with a CNV. This frequency measure facilitates the comparison of catalogs with different methods and resolution. We represented the distribution as a cumulative proportion distribution in Figure 3a. The graphs read as “how much of the total affected genome is called in at more than X% of the population”. The frequency distribution was computed separately for deletions and duplications (and *CNV* in the 1000 Genomes Project catalog). Of note, the 1000 Genomes Project was downsampled to 640 random individuals in order to give comparable frequency curves.

#### Comparison with CNV catalogs from long-read studies

First, the SV catalog from Chaisson et al.^57^ was downloaded from http://eichlerlab.gs.washington.edu/publications/chm1-structural-variat

Recurrent calls were collapsed in both `PopSV` and the 1000 Genomes Project catalogs. `PopSV`’s catalog corresponded to all germline calls in the Twin study, renal cancer dataset and GoNL. The 1000 Genomes Project catalog contained all the deletions, duplications and CNVs, no matter the size or frequency. The analysis was also performed separately on deletions, duplications, low-mappability regions and extremely low-mappability regions. For each comparison, we randomly selected control regions with sizes and overlap with assembly gaps similar to the SVs in Chaisson et al.^57^ (see Selecting control regions). A logistic regression tested the enrichment of CNVs in the Chaisson catalog versus the control regions. The regression was performed on 50 different sampling of the control regions for each comparison. The 50 samplings are represented by the boxplot in Figure 3b. We compared the estimates from the logistic regression. They represent the log odds ratio of a CNV overlapping the catalog from Chaisson. The same analysis was performed using the SV catalog from Pendleton et al.^53^ downloaded from ftp://ftp-trace.ncbi.nlm.nih.gov/giab/ftp/data/NA12878/NA12878_PacBio_MtSinai/NA12878.sorted.vcf.gz.

#### Distance to centromere, telomere and assembly gaps

The centromeres, telomeres and assembly gaps (CTGs) are annotated in the gap track from UCSC^58^. However, some chromosomes were missing telomere annotations. We defined them as the 10 Kbp region at the ends of chromosomes derived from the cytogenetic bands track.

The distance from each variant to the nearest CTG was computed and represented as a cumulative proportion, i.e. the proportion of variants located at a distance *d* or closer to a CTG.

Because this distribution changes with the size of the variants, we sampled random regions in the genome with similar sizes and computed the same distance distribution (see Selecting control regions). Thanks to this null distribution we were able to see if variants were located closer/further to CTG than expected by chance.

#### Selecting control regions

In several analyses we compared the CNVs with control regions. The control regions have the same size distribution as the regions they are derived from (e.g. CNV, annotation). In some analysis we further controlled for the overlap with specific genomic features. For example, we controlled for the overlap with CTGs to avoid selecting control regions in assembly gaps where no CNV or annotation is available. Controlling for the overlap with regions flanking CTGs, we could simply control for the distance to CTGs. We also used this approach to control for the overlap with segmental duplications and investigate patterns independent from this repeat class.

To select control regions, thousands of bases were first randomly chosen in the genome. The distance between each base and the genomic features was then computed. At this point, simulating a region of a specific size and with specific overlap profile can be done by randomly choosing as center one of the bases that fit the profile:

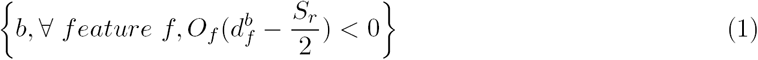

with *O_f_* equals 1 if the original region overlaps with feature *f*, -1 if not; 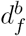 is the distance between base *b* and feature *f*; and *S*_*r*_ is the size of the original region.

For each input region, a control region was selected as described and had by construction the exact same size and overlap profile.

#### Enrichment in genomic features

We tested different genomic features, starting with: genes, exons, low-mappability regions, segmental duplications, satellites, simple repeats and transposable elements. The different satellite families, frequent simple repeat motives, transposable element families were also tested. We overlapped each genomic feature with CNVs and control regions. We then computed the fold change in proportion of regions overlapping a feature, in CNV versus control regions. A pseudo count was added when computing this ratio:

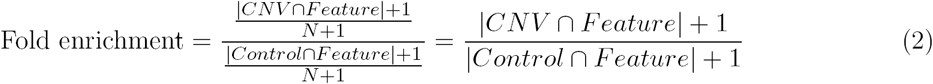

where *N* is the number of CNVs (and control regions).

The fold enrichments were computed separately for each sample using control regions that fitted perfectly the profile of the variants in the sample. To assess the significance of the enrichment, a logistic regression was performed using CNV and control regions. The model to test one feature in one sample was:

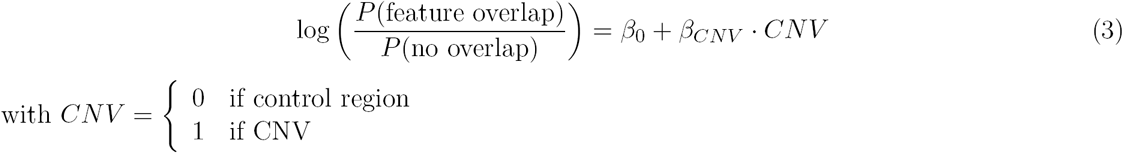

To control for the enrichment in segmental duplication we used control regions with similar overlap profile (see Selecting control regions). We also added a variable representing the overlap with segmental duplication in the model:

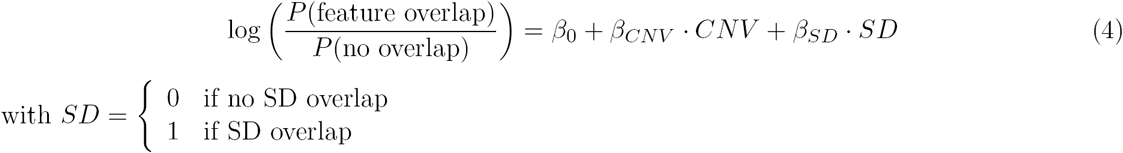

For each feature and cohort we computed the median P-value. When numerous tests were performed (e.g. satellite families, simple repeat motives, transposable element families or subfamilies), the P-values were first corrected for multiple testing using Benjamini-Hochberg procedure.

Finally, we computed the proportion of the region overlapped by the different features (satellites, simple repeats and transposable elements). We compared CNV regions and control regions.

#### Somatic variant definition

Somatic variants were defined as variant in a tumor samples with low overlap with variant in the paired normal sample. In CageKid data, overlapping tumor variant with the ones from the paired normal showed almost only two peaks, at 0 and 100% overlap. A tumor variant was defined as somatic if it overlapped less than 10% of any variant in the paired normal.

